# Kinetochore-associated Mps1 regulates the strength of kinetochore-microtubule attachments via Ndc80 phosphorylation

**DOI:** 10.1101/2021.04.25.441339

**Authors:** Krishna K. Sarangapani, Lori B. Koch, Christian R. Nelson, Charles L. Asbury, Sue Biggins

## Abstract

Dividing cells detect and correct erroneous kinetochore-microtubule attachments during mitosis, thereby avoiding chromosome mis-segregation. Most studies of this process have focused on the Aurora B kinase, which phosphorylates microtubule-binding elements specifically at incorrectly attached kinetochores, promoting their release and providing another chance for proper attachments to form. However, growing evidence suggests additional mechanisms, potentially involving Mps1 kinase, that also underlie error correction. Because these mechanisms overlap *in vivo*, and because both Mps1 and Aurora B function in numerous other vital processes, their contributions to the correction of erroneous kinetochore attachments have been difficult to disentangle. Here we directly examine how Mps1 activity affects kinetochore-microtubule attachments using a reconstitution-based approach that allowed us to separate its effects from Aurora B activity. When endogenous Mps1 that co-purifies with isolated kinetochores is activated *in vitro*, it weakens their attachments to microtubules via phosphorylation of Ndc80, a major microtubule-binding element of the outer kinetochore. Mps1 phosphorylation of Ndc80 appears to contribute to error correction because phospho-deficient Ndc80 mutants exhibit genetic interactions and segregation defects when combined with mutants in an intrinsic error correction pathway. In addition, Mps1 phosphorylation of Ndc80 is stimulated on kinetochores lacking tension. These data suggest that Mps1 provides an additional mechanism for correcting erroneous kinetochore-microtubule attachments, complementing the well-known activity of Aurora B.

## Introduction

The equal segregation of duplicated chromosomes to daughter cells during cell division is fundamental to life. Segregation is mediated by interactions between dynamic microtubules and kinetochores, the megadalton protein complexes that assemble on the centromeres of each chromosome (Monda and Cheeseman, 2018; Musacchio and Desai, 2017). For accurate segregation, the sister kinetochores on each pair of chromatids must make bioriented attachments to microtubules emanating from opposite spindle poles. When linked sister kinetochores achieve biorientation, they come under tension due to pulling forces exerted by the opposing microtubules. However, because kinetochore-microtubule attachments initially form at random, erroneous connections lacking tension are often made. These must be detected and corrected to avoid mis-segregation. Tension appears to help cells distinguish correct from incorrect attachments, because attachments under tension are more stable than those lacking tension *in vivo* and *in vitro* (Akiyoshi et al., 2010; King and Nicklas, 2000; Nicklas and Koch, 1969).

A variety of error correction mechanisms help cells to make proper bioriented attachments. The most well-studied mechanism involves the conserved essential protein kinase Aurora B (Krenn and Musacchio, 2015). When a kinetochore attaches incorrectly, the lack of tension is believed to signal Aurora B kinase to phosphorylate kinetochore proteins, which weakens their grip on the microtubule, causing detachment, and giving the cell another chance to make a proper attachment (Biggins et al., 1999; Dewar et al., 2004; Hauf et al., 2003; Tanaka et al., 2002). When Aurora B is defective, cells are unable to make bioriented attachments (Biggins et al., 1999; Hauf et al., 2003; Tanaka et al., 2002). One of the major Aurora B substrates is Ndc80, a core component of the kinetochore that makes a significant contribution to kinetochore-microtubule coupling (Cheeseman et al., 2006; DeLuca et al., 2006). Ndc80 has two domains that mediate its interaction with the microtubule, a conserved calponin-homology ‘head’ domain and a disordered N-terminal ‘tail’ (Ciferri et al., 2008; Wei et al., 2007). The Ndc80 N-terminal tail contains multiple Aurora B consensus sites and is the site of Aurora B regulation (Akiyoshi et al., 2009; Cheeseman et al., 2006; Ciferri et al., 2008; DeLuca et al., 2006; Guimaraes et al., 2008; Miller et al., 2008; Wei et al., 2007). Additional kinetochore proteins are also phosphorylated by Aurora B to destabilize kinetochore-microtubule attachments but they vary depending on the organism (Cheeseman et al., 2002; Lan et al., 2004; Wordeman et al., 2007).

The Mps1 kinase is another conserved essential kinase implicated in kinetochore biorientation and error correction, independent of its well-studied role in signaling the spindle assembly checkpoint (Hewitt et al., 2010; Jelluma et al., 2008; Jones et al., 2005; Maciejowski et al., 2010; Maure et al., 2007; Santaguida et al., 2010). While there is some evidence that Mps1 regulates Aurora B activity (Jelluma et al., 2010; Saurin et al., 2011; Tighe et al., 2008), significant data suggests it has an independent role in error correction and acts downstream of Aurora B (Hewitt et al., 2010; Maciejowski et al., 2010; Maure et al., 2007; Meyer et al., 2013; Santaguida et al., 2010). Consistent with this interpretation, inhibition of Mps1 does not alter the phosphorylation of many Aurora B substrates or Aurora B localization (Hewitt et al., 2010; Maciejowski et al., 2010; Maure et al., 2007; Santaguida et al., 2010; Tighe et al., 2008). Mps1 localizes to kinetochores by binding to the Ndc80 protein and its activity is highest on kinetochores that have not made proper attachments (Hiruma et al., 2015; Ji et al., 2015; Kemmler et al., 2009; Kuijt et al., 2020). However, its role in regulating kinetochore-microtubule attachments and error correction has not been fully elucidated. In human cells, Mps1 regulates localization of the motor protein CENP-E and the Ska complex, which stabilize proper kinetochore-microtubule attachments (Espeut et al., 2008; Hewitt et al., 2010; Maciejowski et al., 2017; Stucke et al., 2004). In budding yeast, Mps1 is required for localization of the Dam1 complex (Meyer et al., 2018; Meyer et al., 2013), an ortholog of the Ska complex (van Hooff et al., 2017). It also phosphorylates the Ndc80 protein, but this phosphorylation was reportedly involved in spindle checkpoint signaling and not in error correction (Kemmler et al., 2009). Recently, it was reported that Mps1 regulates biorientation via phosphorylation of the Spc105 protein (Benzi et al., 2020). While Mps1 targets many key microtubule-binding kinetochore elements, a unified view of how it participates in error correction remains elusive, in part because it is challenging to disentangle this function from the similar function of Aurora B and the well-established role for Mps1 in checkpoint signaling.

To directly study the effect of Mps1 activity on kinetochore-microtubule attachments, we took advantage of a yeast reconstitution system where the strength of attachment between individual isolated kinetochores and single microtubules can be measured *in vitro* (Akiyoshi et al., 2010; Sarangapani et al., 2013). Using isolated kinetochore particles that copurify with Mps1 kinase but lack other kinase activity (London et al., 2012), we found that Mps1 phosphorylation of Ndc80 directly weakens kinetochore-microtubule attachments, similar to the function of Aurora B. Moreover, phosphorylation of Ndc80 *in vivo* by Mps1 occurs during mitosis when kinetochores are prevented from coming under tension. Cells containing mutations in the Mps1-targeted phosphorylation sites on Ndc80 exhibit genetic interactions and chromosome segregation defects when combined with inhibitory mutations in another, intrinsic error correction pathway (Akiyoshi et al., 2010; Miller et al., 2018; Miller et al., 2016). Taken together, our data suggest that Mps1-mediated phosphorylation of Ndc80 provides an additional mechanism for specifically weakening kinetochore-microtubule attachments that lack tension, complementing the Aurora B and intrinsic error correction pathways, thereby helping to release erroneous attachments and ensure the accuracy of chromosome segregation.

## Results

### Copurifying kinase activity weakens the attachment of isolated kinetochores to microtubules

We previously found that native kinetochore particles purified from budding yeast lack Aurora B kinase activity and that the major copurifying kinase activity is due to Mps1 (London et al., 2012). To test whether the copurifying Mps1 activity directly affects interactions between kinetochores and microtubules, we modified our previously developed approach for measuring the strengths of individual kinetochore-microtubule attachments *in vitro* (Akiyoshi et al., 2010), by adding ATP to activate any copurifying kinase. We isolated kinetochores via anti-Flag immunoprecipitation of the Dsn1 protein (Dsn1-6His-3Flag) (Supplemental Figure S1A). After elution with Flag peptide, we linked the native kinetochore particles to polystyrene microbeads, mixed them with ATP to activate the copurifying kinase, and immediately introduced them into a flow chamber containing dynamic microtubules grown from coverslip-anchored microtubule seeds (Figure 1A). During the time required to seal the slide chamber and mount it onto the laser trap (∼10 min), some kinetochore-decorated beads attached spontaneously to the sides of coverslip-anchored microtubules. The laser trap was used to bring these beads that were laterally attached to microtubules to the plus end tips and then apply gradually increasing force until the kinetochores ruptured from the tips (Figure 1B and 1C). Rupture strengths for many individual attachments were collected in the presence of ATP, or ADP, or without adenosine, until 90 min after sealing the slide. For each population, the median rupture force was calculated and the fraction of attachments that survived up to a given level of force was also plotted. Similar to our previous work (Akiyoshi et al., 2010; Sarangapani et al., 2013), kinetochore particles not exposed to ATP ruptured over a range of forces, with a median strength of 9.8 piconewtons (pN) in the absence of adenosine, or 8.7 pN in the presence of ADP (Figure 1D). However, when ATP was included, the rupture force distribution was shifted to lower values and the median strength was only 5.3 pN, suggesting that phosphorylation by a kinetochore-associated kinase decreased the strength of kinetochore-microtubule attachments. Consistent with this interpretation, if λ-phosphatase was included together with the ATP, then the ATP-dependent weakening was reduced and the kinetochores maintained a median strength of 7.5 pN, similar to untreated and ADP-treated kinetochores (Figure 1D and Supplemental Figure S1B). We monitored the rupture strength as a function of time, which indicated that the ATP-dependent weakening reaction was completed during the ∼10 min slide preparation, with no significant weakening thereafter (Supplemental Figure S1C). Altogether, these data indicate that phosphorylation of native kinetochores by a copurifying kinase activity reduces the strength of their attachments to microtubules.

**Figure 1.**
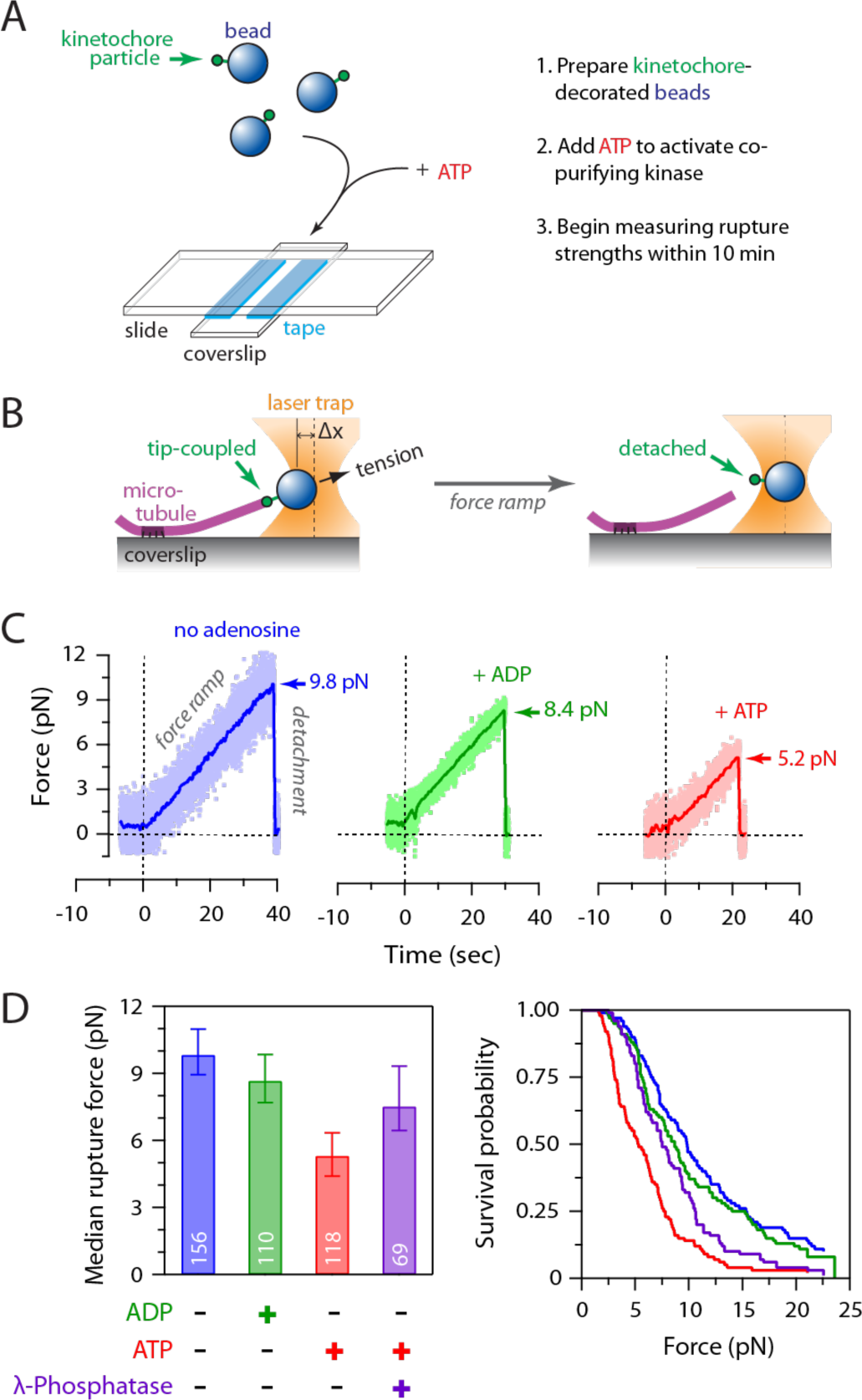
Activity of copurifying kinase weakens attachment of native kinetochores to microtubules. (**A**) Schematic of slide preparation. Kinetochore-deco rated beads were prepared, mixed with ATP (or ADP as a control), free tubulin, and GTP, and then introduced into a flow chamber containing coverslip-anchored microtubul e seeds. (**B**s) Schematic of laser trap assay. After a kinetochore-decorated bead is tip-coupled under low preload (left),the force is gradually increased until detachment (right). Trap measurements commenced ~10 min after mixing kinetochore-decorated beads with ATP (or ADP) and continued for up to 90 min. (**C**) Example traces showing forced detachment under indicated conditions of beads decorated with native kinetochore particles isolated from wild type cells (SBY8253). Dots represent instantaneous force fluctuations. Solid traces show same data after smoothing with a 500-ms sliding boxcar average. Dashed vertical lines mark the start of the 0.25 pN s^-1^ force ramp. Arrows mark rupture force. (**D**) Median rupture strengths (left) and corresponding survival probability distributions (right) for wild type kinetochores (from SBY8253) under indicated conditions. ATP exposure (red) caused significant weakening, which was inhibited by addition of λ-phosphatase (purple). Va lues inside bars ind icate n umbers of events for each condition. Error bars represent ± 95% confiden ce interva ls calculated by bootstrapping (see Supplemental Material).

### Mps1 activity is required for the ATP-dependent weakening of isolated kinetochores

Because Mps1 was the only kinase activity we detected on purified kinetochores (London et al., 2012), we next tested whether Mps1 mediates the ATP-dependent weakening of kinetochore particles using an Mps1 inhibitor called reversine (Santaguida et al., 2010). We first verified that reversine inhibits Mps1 activity on the particles by incubating them with radioactive ATP in the presence or absence of the drug (Figure 2A). Autoradiography revealed phosphorylation on Spc105, Ndc80 and Dsn1 when reversine was omitted (Figure 2A), in agreement with our previous work (London et al., 2012). When reversine was added, total phosphorylation levels on all the substrates decreased by 69%, confirming significant inhibition. We then performed rupture force assays, mixing kinetochore-decorated microbeads with reversine plus ATP, or with the vehicle DMSO plus ATP, ADP, or no adenosine as controls. The median strength of kinetochores measured in the presence of DMSO was 8.2 pN without adenosine and fell to 5.0 pN in the presence of ATP. This loss of strength was completely blocked by reversine (Figure 2B), indicating that the ATP-dependent weakening of isolated kinetochore particles requires Mps1 activity.

**Figure 2.**
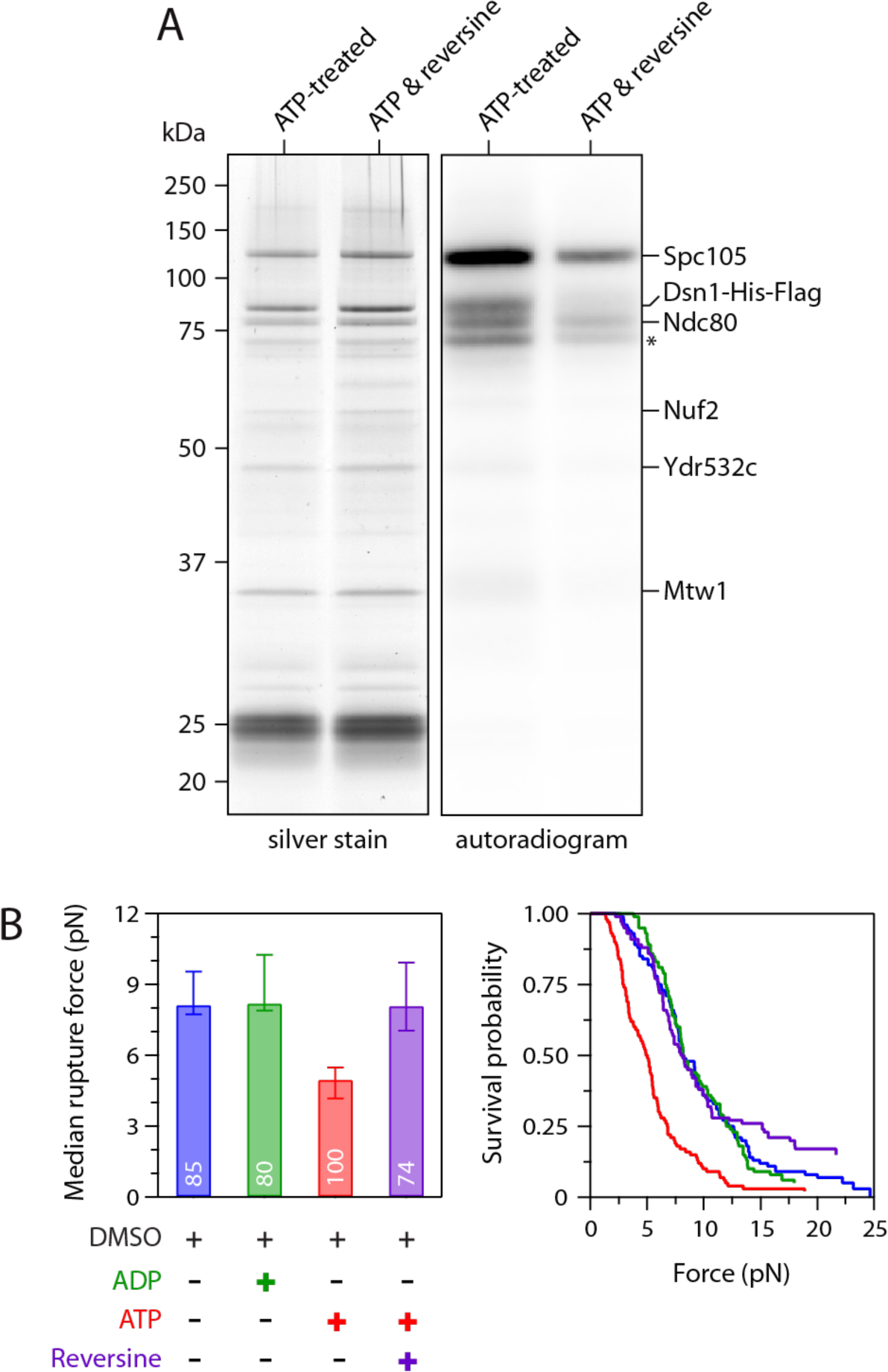
ATP-dependent weakening of native kinetochores requires Mpsl kinase activity. (**A**)The Mpsl inhibitor reversine reduces phosphory­ lation of Spcl 05, Dsn l, and Ndc80 by the copurifying kinase activity. Kinetochores purified by immu noprecipitation of Dsnl -His-Flag from wild type cells (SBY8253) were incu bated with λ-^32^P-labeled ATP, either alone or with 5 µM of reversine, and then visualized by silver stain or autoradiogra­ phy after SOS-PAGE. Asterisk ma rks a n additional u nidentified phosphoprotei n. **(B)** Median rupture strengths (left) and corresponding surviva l probability distributions (right) for wild type kinetochores (from SBY8253) measu red under indicated conditions. The ATP-dependent wea kening (red) was blocked by addition of 5 µM reversine (purple). Values inside bars indicate nu mbers of events for each condition. Error ba rs represent ± 95% confidence intervals calculated by bootstrapping (see Supplemental Material).

### ATP-dependent weakening depends on phosphorylation of Ndc80 not Spc105

We next sought to identify the key substrate(s) whose phosphorylation by Mps1 causes the weakening of reconstituted kinetochore-microtubule attachments. Mps1-mediated phosphorylation of Spc105 is vital for initiating spindle checkpoint signaling (London et al., 2012; Shepperd et al., 2012; Yamagishi et al., 2012), and recent work suggests it also promotes kinetochore biorientation through recruitment of the Bub1 protein (Benzi et al., 2020; Storchova et al., 2011). To test whether Spc105 is the relevant substrate underlying ATP-dependent kinetochore weakening *in vitro*, we purified kinetochores from phospho-deficient *spc105-6A* cells, which carry alanine substitutions to block phosphorylation at all six Mps1 phosphorylation sites (the ‘MELT’ motifs) within the disordered N-terminal region of Spc105 (London et al., 2012) (Figure 3A and Supplemental Figure S2A). In rupture force assays, the median strength of phospho-deficient Spc105-6A kinetochores was 8.3 pN without adenosine and fell to 3.2 pN in the presence of ATP (Figure 3B). We were surprised that the Spc105-6A kinetochores were weaker after ATP addition than their wildtype counterparts, so we analyzed the levels of copurifying Mps1. The phospho-deficient Spc105-6A kinetochores retained more Mps1 than wildtype (Supplemental Figure S2B), potentially explaining their enhanced sensitivity to ATP. While the reason for this higher retention of Mps1 on Spc105-6A kinetochores remains unclear, the rupture strength data nevertheless indicate that Spc105 is not the key Mps1 target underlying ATP-dependent weakening of the kinetochores *in vitro*.

We next focused on the major microtubule binding component of the kinetochore, Ndc80c (Akiyoshi et al., 2010; Cheeseman et al., 2006; DeLuca et al., 2006). Previous work identified a total of fourteen sites on the Ndc80 protein that are phosphorylated by Mps1 (Kemmler et al., 2009) (Figure 3C). However, three of these overlap with known Aurora B phospho-sites (Akiyoshi et al., 2009), so we tested whether the remaining eleven are involved in the ATP-dependent weakening. We purified kinetochore particles from phospho-deficient *ndc80-11A* cells, which carry alanine substitutions at all eleven of the Mps1 phosphorylation sites, to prevent their phosphorylation (Supplemental Figure S2C) (Kemmler et al., 2009). Similar levels of Mps1 copurified with both Ndc80-11A and wild type kinetochores (Supplemental Figure S2D), but the kinase activity had no effect on the rupture strengths of Ndc80-11A kinetochores. The median strength of Ndc80-11A kinetochores was 8.7 pN in the absence of adenosine, similar to wild type kinetochores (Figure 3D). Upon exposure to ATP, the median strength of Ndc80-11A kinetochores remained high, 8.5 pN, and statistically indistinguishable from the strength measured without adenosine (Figure 3D and Supplemental Figure S2E). These data indicate that one or more of the Mps1 phosphorylation sites on Ndc80 are required for the decreased attachment strength.

**Figure 3.**
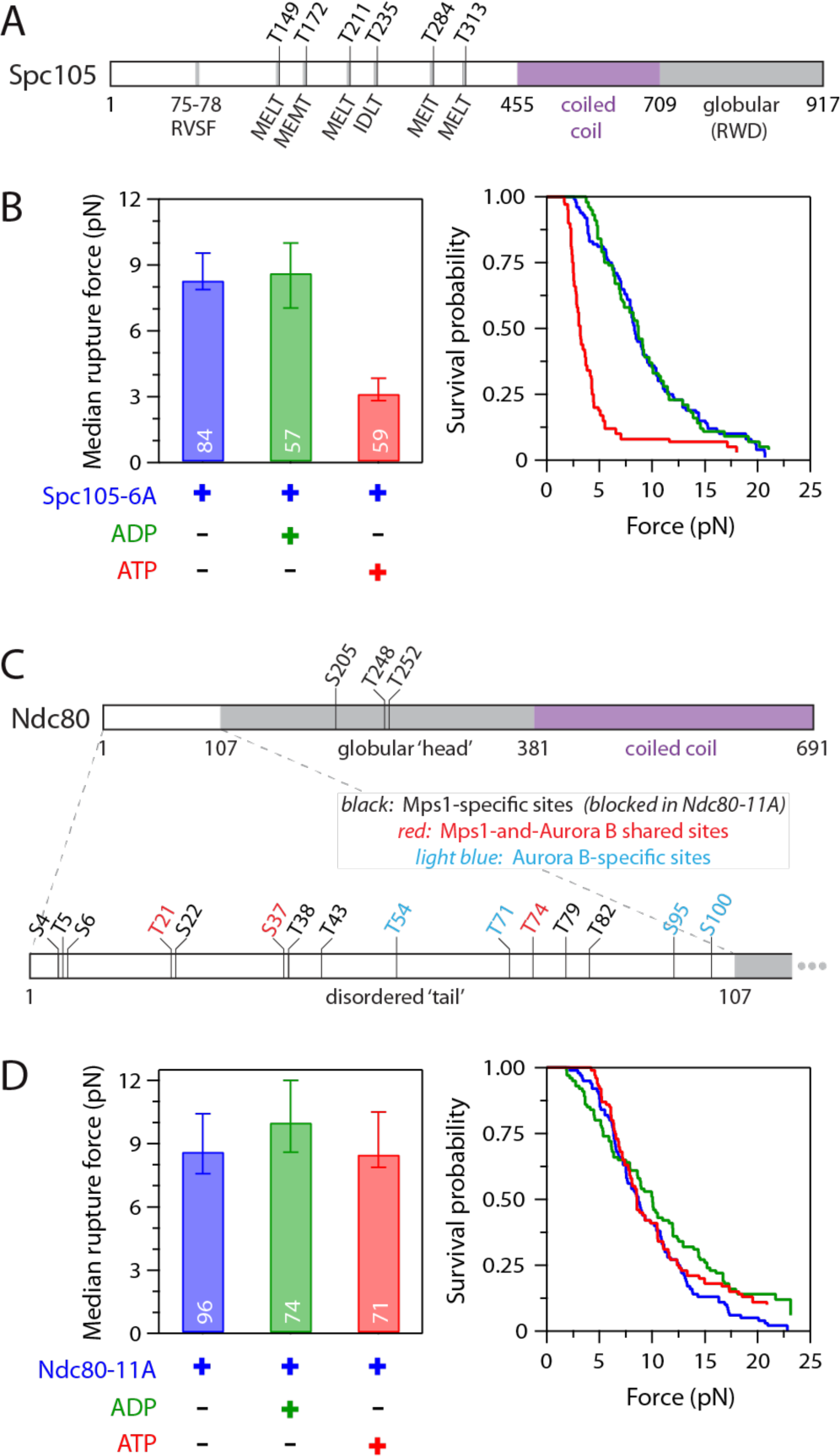
ATP-dependent weakenin g depends on phosphorylation of the Ndc80 tail, not of Spc105. **(A)** Map of six Mpsl phosphorylation target sites (‘MELT’ motifs) within the disordered N-terminus of Spcl 05. **(B)** Median rupture strengths (left) and corresponding survival probability distributions (right) for phospho-deficient Spcl 05-6A kinetochores (from SBYl 0315). These kinetochores were weakened upon exposure to ATP by an amount similar to kinetochores carrying wild type Spcl OS. **(C)** Map of Mpsl and Aurora B phosphorylation target sites within the globular head (top) and disordered N-termina l tail (bottom) of Ndc80. The tail includ es three shared sites targeted by both Mpsl and Aurora B (shown in red), plus eight sites targeted specifica lly by Mpsl (black). (Four additional sites targeted specifically by Aurora B are shown in light blue.) (**D**) Median rupture strengths (left) and correspond ing survival probability distribu­ tions (right) for phospho-defic ient Ndc80-l l A kinetochores (from SBYl 9838), with Ala substitutions blocking all eleven of the Mpsl-specific target sites (i.e., those colored black in C), measured under ind icated conditions. These kinetochores were refra ctory to ATP-dependent weakening. Residue numbers below the maps in A and C demarcate major sequence features. Values inside bars in B and D indicate numbers of events for each condition. Error bars represent ± 95% confidence intervals calculated by bootstrapping (see Supplemental Material).

### Mps1 phosphorylation of Ndc80 is not required for spindle assembly checkpoint signaling

A previous report suggested that Mps1-mediated phosphorylation of Ndc80 regulates the spindle assembly checkpoint and does not affect kinetochore-microtubule attachments or chromosome segregation (Kemmler et al., 2009), a conclusion that seemed inconsistent with our *in vitro* results. We therefore re-examined the phenotypes of mutant *ndc80-14A* cells, with alanine substitutions at all fourteen Mps1 target sites included for consistency with the earlier study. To test whether *ndc80-14A* mutants are defective in the spindle assembly checkpoint, we released wild type and *ndc80-14A* cells from G1 into the microtubule depolymerizing drug nocodazole, which generates unattached kinetochores that normally trigger the spindle assembly checkpoint, and monitored cell cycle progression by analyzing levels of the anaphase inhibitor Pds1/securin (Cohen-Fix et al., 1996). Pds1 levels accumulated and then stabilized in both wild type and *ndc80-14A* cells as they progressed through the cell cycle and then arrested (Figure 4A). The stabilization of Pds1 in both wild type and *ndc80-14A* cells depended on the spindle assembly checkpoint, because it was eliminated in both strains by deletion of the checkpoint gene *MAD2* (Figure 4A). These observations indicate that blocking the phosphorylation of all known Mps1 target sites on the Ndc80 protein does not lead to a defective spindle assembly checkpoint as previously reported (Kemmler et al., 2009). It was also reported that replacing all fourteen Mps1 target residues with phospho-mimetic aspartic acid residues was lethal due to constitutive activation of the spindle assembly checkpoint (Kemmler et al., 2009). To test this, we performed a plasmid shuffle assay using strains expressing a wild-type copy of *NDC80* from a plasmid that also contains the *URA3* gene, which renders cells susceptible to the cytotoxic drug 5-Fluoroorotic acid (5-FOA) (Boeke et al., 1987). Cells carrying a functional chromosomal copy of *NDC80* can spontaneously lose the *NDC80-URA3* plasmid and are therefore able to grow on media containing 5-FOA, whereas cells with a non-functional chromosomal allele require the plasmid for viability and are killed by 5-FOA. Using this assay, we confirmed that *ndc80-14D* is recessive lethal (Figure 4B). However, in contrast to the previous report (Kemmler et al., 2009), deleting the spindle assembly checkpoint gene *MAD2* did not rescue this lethality (Figure 4B). Notably, spurious colonies appeared after the *ndc80-14D* mutants were grown for longer times, suggesting the existence of spontaneous suppressor mutations (Supplemental Figure S3). While our data indicate that mutation of the previously identified Mps1 phosphorylation sites on Ndc80 does not cause defects in spindle assembly checkpoint signaling, they confirm the previously reported lethality of *ndc80-14D* (Kemmler et al., 2009). Therefore, one or more of the Mps1 phosphorylation sites have a critical cellular function.

**Figure 4.**
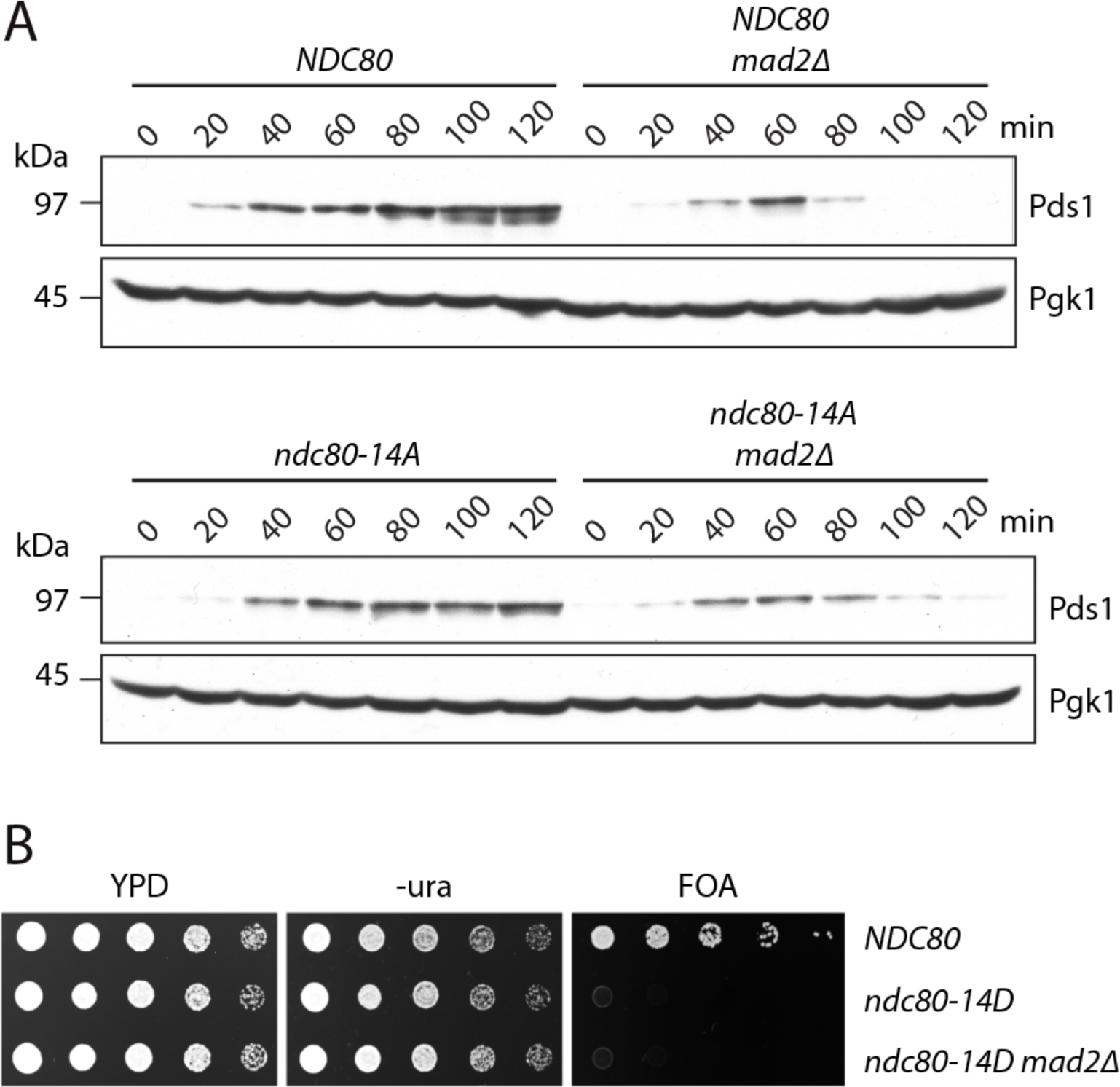
Mpsl phosphoryl ation of Ndc80 is not req uired for spindle assembly checkpoint signaling. **(A)** Cells carrying phospho-deficient *ndc80-14A* maintain a spindle assembly checkpoint arrest in response to nocodazole. The indicated strains (SBYl 7648, SBYl 7624, SBYl 7807, SBYl 7895) were i nitially arrested in G 1 using α-factor for 3 h. α-factor was then washed out and the cells were resuspended in media with 10 µg/ml nocodazole. Samples taken at indicated timepoints were analyzed for the anaphase in hibitor Pdsl /securin (and Pgkl as a loading control) by immunoblotting. **(B)** The phospho-mimetic *ndc80-14D* allele is recessive lethal and its letha lity is not rescued by deletion of the checkpoint gene *MAD2*. The ind icated strains (SBYl 5149, SBY9453, SBYl 5087) were tested using a plasmid shuffle assay, where growth on 5′fluoruracil (FOA) plates ind icates via bility in the absence of a covering copy of wild type *NDC80*. Five-fold serial d ilutions were plated onto YPD, -ura, or FOA plates and grown at 23 °C.

### Weakening occurs via phosphorylation of Mps1-specific targets in N-terminal tail of Ndc80

Our observation that Mps1-mediated phosphorylation of Ndc80 weakens reconstituted kinetochore-microtubule attachments is similar to the well-documented function of Aurora B-mediated phosphorylation of Ndc80, which inhibits kinetochore-microtubule attachments *in vivo* and *in vitro* (Cheeseman et al., 2006; Ciferri et al., 2008; DeLuca et al., 2006; Sarangapani et al., 2013; Wei et al., 2007). Eight of the eleven Mps1-specific target sites on Ndc80 fall within the disordered N-terminal tail domain, which is also where the key Aurora B target sites are located (Figure 3C). To analyze the contribution of the Mps1-specific sites within the Ndc80 tail (Figure 5A), we generated a mutant with alanine substitutions at just these eight sites (Nc80-8A). We initially included an epitope tag (-3HA) to allow direct immunoprecipitation of the phospho-deficient Ndc80-8A, or of wild type Ndc80 as a control. Mps1 copurified with wild type Ndc80, as previously demonstrated (Kemmler et al., 2009), and we found that a similar level of Mps1 copurified with Ndc80-8A (Figure 5B). After incubating both immunoprecipitations with radioactive ATP, autoradiography showed 49% less phosphorylation on Ndc80-8A relative to wild type (Figure 5B), confirming that Mps1 phosphorylates Ndc80 at one or more of the eight Mps1-specific target sites within its N-terminal tail domain.

**Figure 5.**
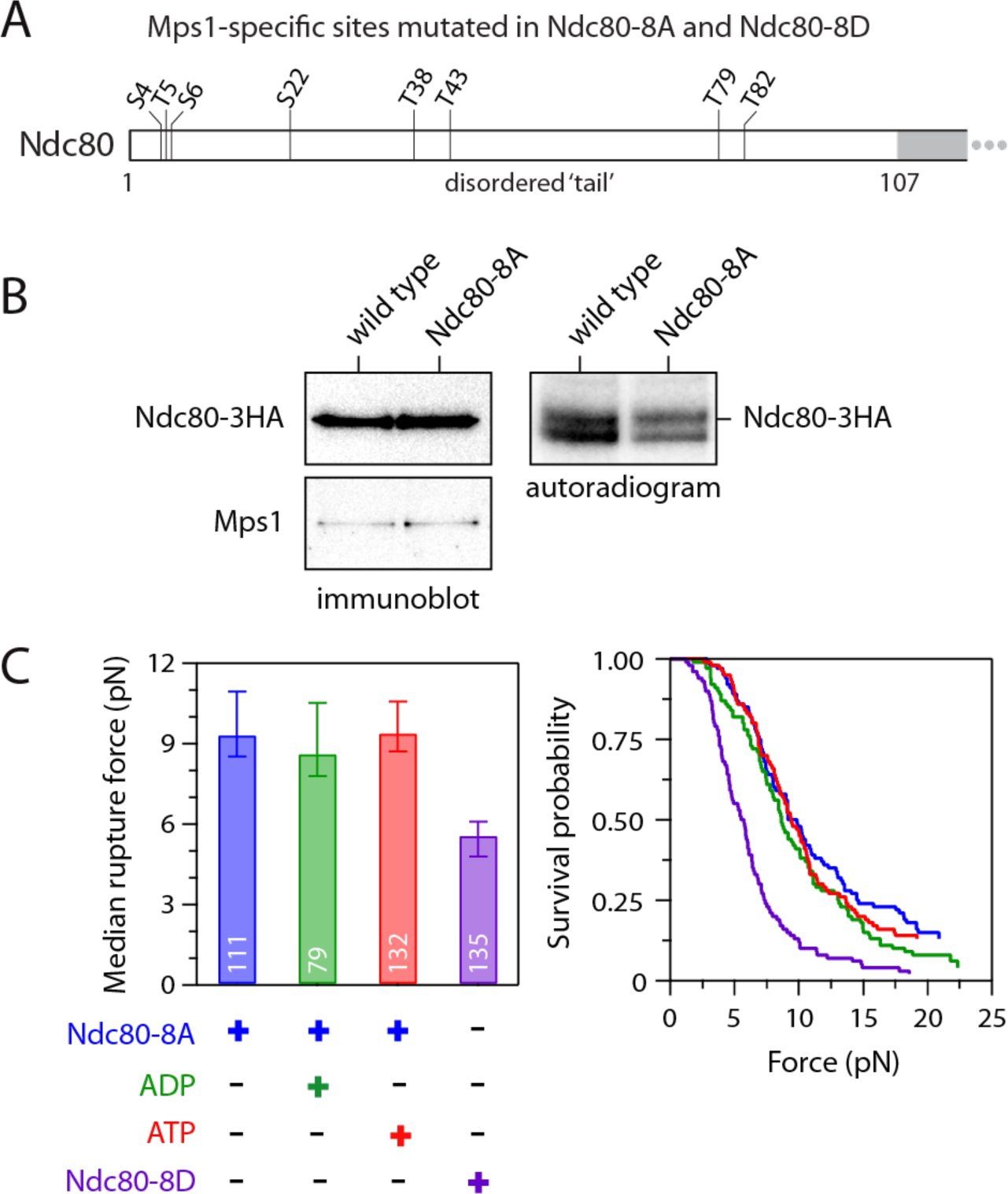
ATP-dependent weakening depends on phosphorylation of Mpsl-specific target sites in the Ndc80 tail. **(A)** Map of Mpsl-specific target sites within the N-terminal tail of Ndc80. All eight sites were mutated to Ala in phospho-deficient Ndc80-8A and to Asp in phospho-mimetic Ndc80-8D. **(B)** *In vitro* phosphorylation of Ndc80-8A was reduced relative to wild type Ndc80,even though similar levels of Mpsl were co-purified. Wild type Ndc80 and phospho-defic ient Ndc80-8A were purified by immunoprecipitation (from SBY20062 and SBY20063), incubated with ^32^P-λ-labeled ATP,and then visualized by immunoblotting (left) or autoradiography (right) after SOS-PAGE. **(B)** Median rupture strengths (left) and corresponding survival probability distributions (right) for phospho-deficient Ndc80-8A and phospho-mi­metic Ndc80-8D kinetochores (purified from SBY19855 and SBY19877, respectively) measured under indicated conditions. The Ndc80-8A kinetochores were refractory to ATP-dependent weakening (red), whereas Ndc80-8D kinetochores were constitutively weak (purple). Values inside bars indicate numbers of events for each condition. Error bars represent ± 95% confidence intervals calculated by bootstrapping (see Supplemental Material).

To test whether these Mps1-specific sites in the Ndc80 tail are involved in ATP-dependent weakening of kinetochore-microtubule attachments, we isolated kinetochores from strains containing wild type or Ndc80-8A protein (via anti-Flag-based immunoprecipitation of Dsn1-6His-3Flag). Based on silver-staining after SDS-PAGE, the composition of the purified Ndc80-8A kinetochore was similar to wild type (Supplemental Figure S4A). In the rupture force assay, the Ndc80-8A kinetochores had a median strength of 9.4 pN when no adenosine was included, and a strength of 8.6 pN in the presence of ADP, values similar to wild type kinetochores (Figure 5C). However, unlike wild type kinetochores, the Ndc80-8A kinetochores were unaffected by exposure to ATP, maintaining a high median rupture strength of 9.4 pN (Figure 5C). To further test the importance of the Mps1-specific tail sites, we also purified Ndc80-8D kinetochores carrying aspartic acid substitutions at all eight sites, to mimic their phosphorylation. The phospho-mimetic Ndc80-8D kinetochores had a composition similar to wild type kinetochores (Supplemental Figure S4A), but their rupture strength was constitutively low, with a median rupture strength of only 4.8 pN in the absence of adenosine (Figure 5C). Altogether, these observations suggest that Mps1 phosphorylation of the Ndc80 tail is necessary and sufficient for directly weakening the isolated kinetochores upon exposure to ATP.

### Mps1 phosphorylation of Ndc80 regulates kinetochore function *in vivo*

To determine whether Mps1 phosphorylates Ndc80 *in vivo*, we set out to generate an antibody that recognizes an Mps1 site. Although we were not able to produce one against a site specific to Mps1 phosphorylation, we succeeded in generating a polyclonal antibody that recognizes Ndc80 carrying a phosphate modification at Thr-74, a site that is phosphorylated by both Ipl1 and Mps1 (Akiyoshi et al., 2009; Kemmler et al., 2009). On immunoblots of purified kinetochore complexes, this Ndc80-T74P antibody detected a signal corresponding to the Ndc80 protein that was diminished when the kinetochores were treated with λ-phosphatase (Figure 6A), confirming the phospho-specificity of the antibody. To check whether Mps1 contributes to the phosphorylation of this site, we purified kinetochores from mutant *mps1-1* cells, in which Mps1 is specifically inactivated when the cells are grown at a non-permissive temperature (Figure 6B). The Ndc80 signal detected in wild type kinetochore particles by the Ndc80-T74P antibody was reduced when mutant *mps1-1* kinetochores were probed. To confirm that Aurora B also phosphorylates this residue, we purified kinetochores from *ipl1-321* cells that lack Aurora kinase activity and also found a reduction in phosphorylation of T74 on Ndc80 (Biggins et al., 1999) (Figure 6C). Taken together, these data show that both Mps1 and Aurora B contribute to Ndc80-T74 phosphorylation on kinetochores *in vivo*.

**Figure 6.**
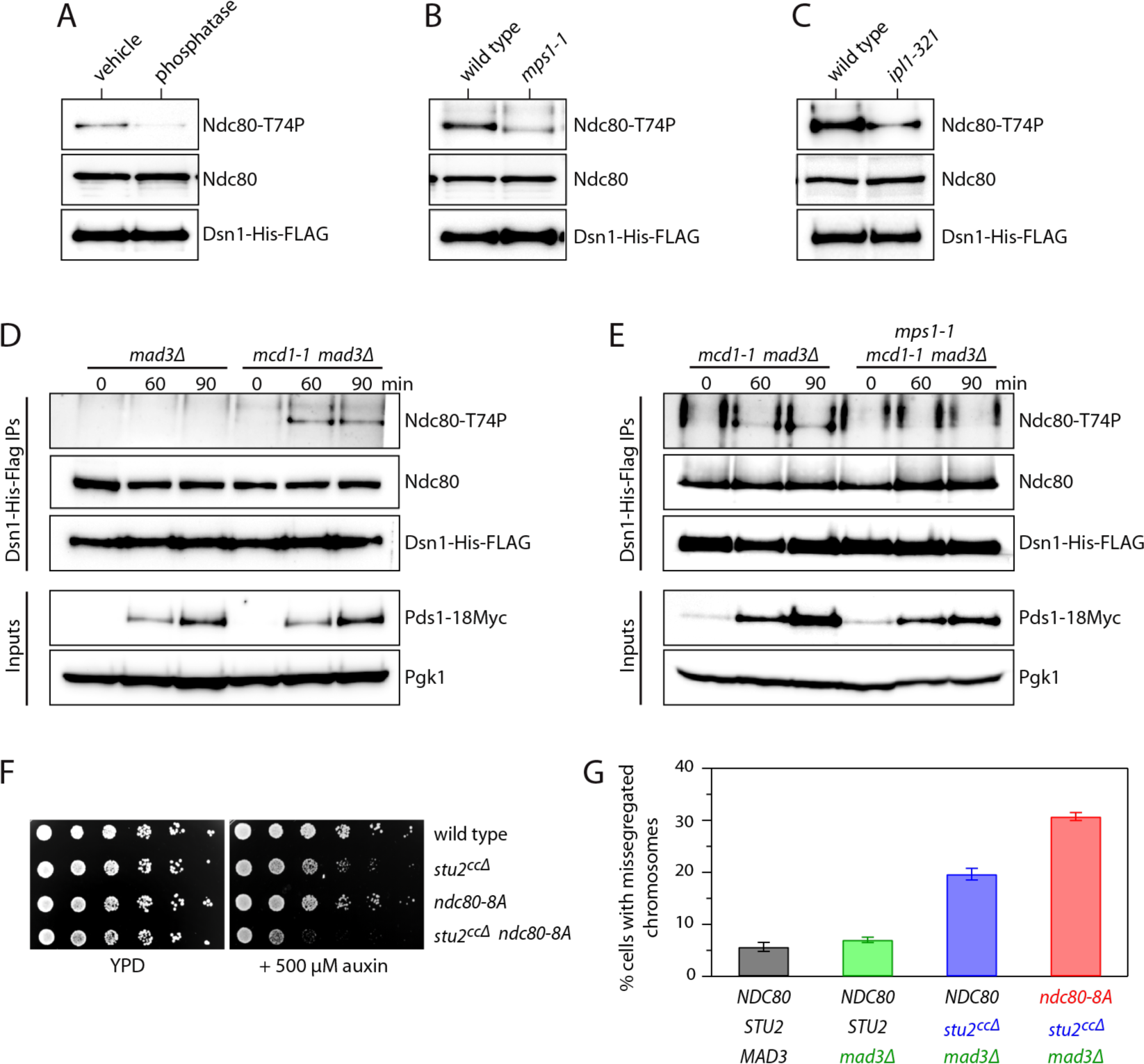
Mpsl phosphorylation of Ndc80 regulates kinetochore function *in vivo.* **(A)** Kinetochores purified from wild type cells (SBY8253) were treated with or without A-phosphatase at 30 °(for 30 min. Phosphorylation of Ndc80 at Thr-74 was visualized by immunoblotting. **(B)** Kinetochores were purified from wild type (SBY8253) or *mpsl -1* cells (SBY8726) after a 3 hr 37 ° C temperature shift. Phosphorylation of Ndc80 at Thr-74 was visualized by immunoblotting. (C) Kinetochores were purified from wild type (SBY8253) or *ip/1-321* cells (SBY8721) after a 3 hr 37 °C temperature shift. Phosphorylation of Ndc80 at Thr-74 was visualized by immunoblotting. **(D)** Phosphorylation of Ndc80 on kinetochores at Thr-74 was analyzed 30, 60, and 90 min after G1 release in *mad3*Δ cells (SBY20362) and *mad3Δ mcd l-1* cells (SBY20361) via immunoblotting. Cells were arrested in α-factor, released at 37 °C, and kinetochore purifications were performed at indicated timepoints. Pdsl-18Myc immunoblotting was used to indicate mitotic timing. **(E)** Phosphorylation of Ndc80 on kinetochores at Thr-74 was analyzed as in (D) in *mad3Δ mcd l -1* cells (SBY20361) and *mad3Δ mcd l-1 mps i-1* cells (SBY20622). **(F)** Wild type cells *(Ndc80-3HA;* SBY20062), *stu2-cc*Δ cells expressing a *stu2-AID* system *(Ndc80-3HA;* SBY20170), *ndc80-8A* cells *(ndc80-8A-3HA;* SBY20063), and *ndc80-8A stu2-cc*Δ cells expressing a *stu2-AID* system *(ndc80-8A-3HA;* SBY20171) were serially diluted and spotted on YPD or YPD + 500 µM auxin plates at 30°C. **(G)** Segregation of chromosome 8 was analyzed in *stu2-AID* cells expressing *Stu2-3V5* (SBYl 7527), *Stu2-3V5 mad3*Δ (SBYl 7668), *stu2-ccΔ mad3Δ Ndc80-3HA* (SBY20210), or *stu2-ccΔ mad3Δ ndc80-8A-3HA* (SBY20211). Segregation was scored in large-budded cells 2 hr after α-factor release.

A lack of tension on erroneous kinetochore-microtubule attachments is thought to promote their release by triggering Aurora B-mediated kinetochore phosphorylation (Biggins and Murray, 2001; Nicklas and Koch, 1969; Stern and Murray, 2001; Tanaka et al., 2002). To test whether a lack of tension can also trigger Mps1-mediated kinetochore phosphorylation *in vivo*, we analyzed the phosphorylation of Ndc80-T74 on kinetochores in a mutant *mcd1-1* strain, which cannot generate tension on its kinetochores due to a defect in sister chromatid cohesion (Stern and Murray, 2001; Tanaka et al., 2000). Wild type and *mcd1-1* mutant cultures were released from G1 to the non-permissive temperature and kinetochores were purified at various time points as the cells progressed through the cell cycle. The cells had *MAD3* deleted to ensure equivalent progression through the cell cycle, which we verified by analyzing Pds1 levels in the lysates (Figure 6D). Immunoblotting confirmed that kinetochores purified from either wild type or *mcd1-1* cells contained similar amounts of Ndc80 protein. In the *mad3*Δ background, we detected weak Ndc80-T74 phosphorylation on wild type kinetochores at time points corresponding to mitosis. However, kinetochores purified from the cohesion-deficient *mcd1-1* cells showed an enrichment of phosphorylation at Ndc80-T74 (Figure 6D), indicating that this phosphorylation is enhanced *in vivo* under conditions where kinetochores lack tension. To test whether Mps1 contributes to this phosphorylation, we repeated the experiment and compared *mcd1-1 mad3*Δ cells to *mcd1-1 mad3Δ mps1-1* cells. Strikingly, the phosphorylation that occurs in the cohesion mutant was reduced in the absence of Mps1 activity (Figure 6E), suggesting that Mps1 contributes to Ndc80 phosphorylation *in vivo* in response to tension defects.

While Aurora B and Mps1 both regulate the N-terminus of Ndc80, mutating their phosphorylation sites in the Ndc80 tail is not lethal (Figure 6F and (Akiyoshi et al., 2009)), consistent with additional pathways contributing to error correction. One such mechanism is the direct stabilization of kinetochore-microtubule attachments by tension (Akiyoshi et al., 2010), an intrinsic property of kinetochores that requires kinetochore-bound Stu2 (Miller et al., 2018; Miller et al., 2016). We previously found that mutants in Aurora B exhibit genetic interactions with *stu2^ccΔ^*, an allele of Stu2 that specifically causes defects in kinetochore biorientation (Miller et al., 2018). We therefore tested whether the Mps1-dependent error correction pathway we identified also exhibits growth defects in combination with *stu2^ccΔ^*. Although the *ndc80-8A* cells carrying phospho-blocking substitutions at Mps1-specific target sites are viable, combining *ndc80-8A* with *stu2^ccΔ^* led to a growth defect (Figure 6F). Thus we analyzed chromosome segregation in these cells by generating *ndc80-8A stu2^ccΔ^* strains with a fluorescent marker on a single chromosome and with the endogenous wild type *STU2* gene under control of a conditional auxin-inducible degron (*stu2-AID*) to maintain viability. Because *stu2^ccΔ^* causes a spindle assembly checkpoint-dependent arrest, we also deleted the *MAD3* gene to enable the cells to progress to anaphase. After arresting the cells in G1, we released them into growth media with auxin to repress the endogenous *stu2-AID* gene and then analyzed chromosome segregation in anaphase. Consistent with our prior work (Miller et al., 2018), chromosome mis-segregation was increased in *stu2^ccΔ^ mad3*Δ cells relative to *mad3*Δ control cells (6.5% in *STU2^WT^ mad3*Δ cells versus 19.4% in *stu2^ccΔ^ mad3*Δ; Figure 6G). The addition of *ndc80-8A* strongly exacerbated the chromosome mis-segregation defect (30.3%), indicating that Mps1 phosphorylation of Ndc80 contributes to accurate chromosome segregation.

## Discussion

To study the effects of kinetochore-associated Mps1 on kinetochore-microtubule attachments independently of other cellular kinases or pathways, we devised a reconstitution approach where the endogenous Mps1 that copurifies with isolated kinetochores is activated via exposure to ATP immediately before laser trap strength measurements. Our data show that Mps1-mediated phosphorylation of the N-terminal tail of Ndc80 causes a direct weakening of the kinetochore-microtubule interface *in vitro*. This function was previously difficult to identify *in vivo* due to the presence of Aurora B that can also phosphorylate Ndc80. Our *in vitro* approach allowed us to separate their functions. We further show that Mps1 phosphorylation of Ndc80 occurs *in vivo*, and is enhanced when kinetochores lack tension due to defective sister chromatid cohesion. Moreover, specifically blocking this Mps1-mediated phosphorylation exacerbates chromosome mis-segregation when combined with a *stu2* mutant defective in another kinase-independent, intrinsic error correction pathway (Miller et al., 2018; Miller et al., 2016). Together, these observations suggest that Mps1 contributes directly to the release of erroneous kinetochore-microtubule attachments.

A previous report identified a total of fourteen Mps1 target sites on Ndc80 and suggested that their phosphorylation was important for regulating the spindle assembly checkpoint without affecting kinetochore-microtubule attachments (Kemmler et al., 2009). While we confirmed the lethality of the phospho-mimetic *ndc80-14D* mutant, its lethality was not suppressed in our hands when the checkpoint was eliminated, nor was our phospho-deficient *ndc80-14A* strain defective in checkpoint signaling as reported (Kemmler et al., 2009). These differences in our findings may be due to different strain backgrounds or the possibility that suppressors can arise more easily in the absence of the checkpoint. The *ndc80-8D* strain with phospho-mimetic substitutions at only the eight Mps1-specific target sites within the Ndc80 tail produced constitutively weak kinetochores but nevertheless remained viable. Its viability indicates that the lethality of *ndc80-14D* probably depends on phosphorylation of Mps1 sites outside the N-terminal tail whose function will be important to identify in the future.

Phosphorylation at one or more of the eight Mps1-specific target sites within the Ndc80 tail is necessary and sufficient for Mps1-mediated weakening of reconstituted kinetochore-microtubule attachments. While other yeast kinetochore proteins have been identified as substrates of Mps1 (Benzi et al., 2020; London and Biggins, 2014a; London and Biggins, 2014b; Shimogawa et al., 2006), Spc105 phosphorylation does not appear to directly regulate kinetochore attachment strength. The Ndc80 tail is also a major target of regulation by Aurora B kinase, which in yeast phosphorylates four distinct Aurora B-specific phospho-sites, plus three shared sites phosphorylated by both Mps1 and Aurora B. Phosphorylation of the human Ndc80 (Hec1) tail by Cdk1 kinase was also identified recently and implicated in the correction of erroneous kinetochore-microtubule attachments (Kucharski et al., 2021). Aurora A also phosphorylates a site in the Ndc80 tail during mitosis (DeLuca et al., 2018). Why multiple different kinases phosphorylate a single domain within Ndc80 is not yet clear, but this convergence might allow weakening of kinetochore attachments in response to different kinds of error signals, potentially with different cell cycle timing. At least three other outer kinetochore components are regulated by both Aurora B and Mps1: the yeast Dam1 complex (Cheeseman et al., 2002; Shimogawa et al., 2006), its functional metazoan counterpart the Ska complex (Maciejowski et al., 2017; Redli et al., 2016), and the CENP-E motor protein (Espeut et al., 2008; Kim et al., 2010). Phospho-proteomic analysis also suggests that Ndc80 might be an Mps1 target in human cells (Maciejowski et al., 2017). Thus, outer kinetochore function could be regulated convergently by both kinases in multiple ways and across species.

Mps1-mediated phosphorylation of the Ndc80 tail causes a weakening of kinetochores similar to that caused by phospho-mimetic substitutions at all seven Aurora B target sites within the tail (Sarangapani et al., 2013). This similarity is consistent with the simple view that Ndc80 tail phosphorylation provides rheostat-like control of kinetochore attachment strength (Zaytsev et al., 2015), with each phosphorylation event contributing equally and the resultant strength varying in proportion to the total number of unphosphorylated tail sites. However, a more complex view is suggested by the recent finding that some sites in the Ndc80 tail are dephosphorylated at metaphase while others remain phosphorylated throughout the cell cycle (DeLuca, 2017; Kucharski et al., 2021). Evidently not all phosphorylation sites are redundant, and therefore some might make differential contributions to chromosome segregation. In the future, it will be important to determine whether Aurora B and Mps1 act sequentially or simultaneously and whether any specific phosphorylation sites are more important for the regulation of kinetochore attachment strength to better understand why cells utilize multiple kinases to regulate the same sites (DeLuca et al., 2018; Kucharski et al., 2021; Zaytsev et al., 2015).

Although both Mps1 and Aurora B kinase can directly weaken kinetochores, mutating either one causes significant biorientation defects *in vivo*, which indicates that the two cannot fully compensate for one another to support this function. Such a requirement for both kinases *in vivo*, despite their similar direct effects *in vitro*, could arise because they each respond to different types of attachment errors or with different timing. Presumably it also reflects additional indirect effects that occur in cells when these kinases are mutated. If Aurora B is defective, for example, the opposing phosphatase PP1 prematurely localizes to kinetochores (Rosenberg et al., 2011), potentially causing dephosphorylation of Ndc80 and counteracting the role of Mps1 in error correction. Conversely, reducing Mps1 activity likely decreases Aurora B activity on kinetochores in budding yeast via regulation of the Bub1-shugoshin pathway (Storchova et al., 2011; Verzijlbergen et al., 2014; Yahya et al., 2020). Mps1 recruits the Bub1 kinase to kinetochores via phosphorylation of the Spc105 kinetochore protein (London et al., 2012; Peplowska et al., 2014; Shepperd et al., 2012; Yamagishi et al., 2012), which leads to Bub1-mediated phosphorylation of H2A and the subsequent recruitment of Sgo1 and Aurora B (Kawashima et al., 2010; Storchova et al., 2011; Verzijlbergen et al., 2014). Thus reducing Mps1 activity could indirectly reduce Aurora B activity at kinetochores to a level that is insufficient for error correction. The Bub1-Sgo1 pathway also contributes to kinetochore biorientation (Fernius and Hardwick, 2007; Peplowska et al., 2014), and it was recently reported that Mps1 phosphorylation of Spc105 is a key Mps1 target for kinetochore biorientation *in vivo* (Benzi et al., 2020). We found that Spc105 phosphorylation does not contribute to the Mps1-dependent weakening of reconstituted kinetochore-microtubule attachments *in vitro*. This result is consistent with our purified kinetochores lacking the Bub1-Sgo1 pathway and suggests that Mps1 plays multiple roles in achieving proper kinetochore-microtubule attachments *in vivo*. In the future, it will be important to understand the interplay between these pathways.

In summary, cells appear to rely on multiple kinetochore kinases as well as intrinsic kinase-independent mechanisms to avoid erroneous kinetochore-microtubule attachments during mitosis. While this complexity reflects sophisticated regulation, it also poses a major challenge for distinguishing direct contributions of specific kinases and substrates from indirect effects and, more generally, for developing a complete understanding of mitotic error correction. The combination of *in vitro* reconstitution with *in vivo* analysis that we used here to uncover a direct regulation of kinetochore attachment strength by Mps1 should be useful in the future for further dissection of the vital processes by which dividing cells ensure the accuracy of chromosome segregation.

## Acknowledgements

We are grateful for feedback and critical reading of the manuscript from members of the Biggins and Asbury Labs, as well as the Seattle Mitosis Group. We are grateful to Arshad Desai and the Lechner Lab for contributing reagents. L.B.K. was supported by a National Science Foundation Graduate Research Fellowship (DGE-1256082). This work was supported by a Packard Fellowship 2006-30521 (to C.L.A.), NIH grants R01GM079373, P01GM105537, R35GM134842 (to C.L.A) and R01GM064386 (to S.B.), and by the Genomics and Scientific Imaging Shared Resources of the Fred Hutch/University of Washington Cancer Consortium (P30 CA015704). S.B. is an investigator of the Howard Hughes Medical Institute.

## Materials and Methods

### Strain Construction and yeast techniques

#### Yeast strains and plasmids

*Saccharomyces cerevisiae* strains used in this study are derivatives of SBY3 (W303) and described in Supplemental Table 1. Standard media and microbial techniques were used and yeast strains were constructed via standard genetic techniques (Rose et al., 1990). Construction of *stu2-3HA-IAA7* and *stu2^Δcc^* (*pSTU2-stu2(Δ658-761::GDGAGL^linker^)-3V5*) are described in (Miller et al., 2018; Miller et al., 2016), *DSN1-6His-3Flag* is described in (Akiyoshi et al., 2010), *spc105-6A* is described in (London et al., 2012), *CEN8::lacO:TRP1* and *pCUP-GFP-LacI*12 are described in (Biggins et al., 1999; Miller et al., 2018; Straight et al., 1996). Yeast strains with *ndc80-14A-3HA* were made by transformation with plasmid SB1848 (pSK1039, *pNDC80-ndc80-14A-3HA:KANMX*), a kind gift from Johannes Lechner, or by crossing to derivative strains. Similarly, *ndc80-14D-3HA* yeast were made by transformation of plasmid SB1577 (pSK981, *pNDC80-ndc80-14D-3HA:KANMX*), a kind gift from Johannes Lechner, or by crossing to derivative strains. Strains containing *ndc80-11A-3HA* were made by transformation of plasmid SB3131, which was made by reverting T21 S37 and T74 in the *ndc80-14A* plasmid SB1848 (pSK1039, Lechner Lab) via overlapping primer mutagenesis. *ndc80-8A-3HA* and *ndc80-8D-3HA* were made via standard cloning techniques from SB2412 (*Ndc80-3HA*). Briefly, pSB2412 and gBlocks (IDT Technologies) containing *ndc80-8A* (SB7002) or *ndc80-8D* (SB7001) were digested with BglII and BstEII and ligated using T4 DNA Ligase (NEB). Plasmids were fully sequenced. All plasmids are described in Supplemental Table 2 and primers in Supplemental Table 3.

#### Auxin inducible degradation

The auxin inducible degradation system was used as previously described (Miller et al., 2016). Cells expressing a C-terminal fusion of an auxin responsive protein (IAA7) in the presence of *TIR1*, which is required for auxin induced degradation, were treated with 500μM IAA (indole-3-acetic acid dissolved in DMSO; Sigma) to induce degradation of the desired AID-tagged target protein. For Fig 6G, auxin was added immediately after cells were released from alphα-factor.

#### Serial dilution assay

The indicated yeast strains were grown overnight in YPD (2% glucose) medium. The next day, the cells were diluted to OD_600_ ∼1.0. A serial dilution (1:5) series was made in a 96-well plate and cells were spotted onto YPD or YPD + 500μM auxin (indole-3-acetic acid dissolved in DMSO; Sigma) plates. Plates were incubated for 1-3 days at 30 °C unless otherwise indicated.

#### Chromosome segregation and time course assays

Cells were grown at 23 °C in YPDA medium (2% glucose, 0.02% adenine). For the spindle assembly checkpoint assays, exponentially growing cells were arrested in G1 with 1 μg/ml alpha factor for 3-4 hours, washed 3 times, and then resuspended in medium lacking pheromone but containing 10ug/ml nocodazole. Samples were collected at the indicated times. 1 μg/ml alpha factor was added to the cultures 40-50 minutes after the G1 release to prevent cells from entering a second cell cycle.

To analyze chromosome segregation, exponentially growing *MATa* cells containing a LacI-GFP fusion and 256 *lacO* sequences integrated proximal to *CEN8* were arrested in G1 with 1μg/mL alphα-factor. After 2.5 hours, cells were washed and released into medium lacking pheromone, but containing 500μM IAA. Alphα-factor was re-added ∼75 minutes after release to prevent entry into the subsequent cell-cycle. 120 minutes after G1 release, aliquots of cells were fixed with 3.7% formaldehyde in 100mM phosphate buffer (pH 6.4) for 10 min. Cells were then washed once with 100mM phosphate buffer (pH6.4), and then permeabilized and stained with DAPI by resuspending in 1.2M Sorbitol/1% Triton X-100/100mM phosphate buffer (pH 7.5) containing 1ug/mL DAPI (Molecular Probes) for 5 min. Cells were then resuspended in the same buffer lacking DAPI and applied to a coverslip treated with 0.5mg/mL concanavalin A. Cells were imaged on a Deltavision Ultra deconvolution high-resolution microscope equipped with a 100x/1.4 PlanApo N oil-immersion objective (Olympus) with a 16-bit sCMOS detector. Cells were imaged in Z-stacks through the entire cell using 0.2 μM steps. All images were deconvolved using standard settings and analyzed softWorX 7.2.1(GE).

To analyze phosphorylation of Ndc80 during the cell cycle, cells were initially grown at 23°C in YPD medium (2% glucose). Exponentially growing *MATa* cells were arrested in G1 with 1μg/mL alphα-factor. After 2.5 hours, cells were washed and released into medium lacking pheromone and shifted to 37°C to inactivate *mcd1-1*. 100mL aliquots of cells were taken at 0, 60, and 90 minutes. Kinetochore purification assays were performed as described below for each timepoint and phosphorylation of Ndc80 threonine-74 was analyzed via immunoblot.

### Protein biochemistry

#### Kinetochore purification assay

Native kinetochore particles were purified essentially as previously described (Akiyoshi et al., 2010; Miller et al., 2016). Kinetochores were purified from asynchronously grown *S. cerevisiae* cells grown in YPD medium (2% glucose) unless otherwise noted in the text. Protein lysates were prepared using a Freezer Mill (SPEX SamplePrep) submerged in liquid nitrogen. Lysed cells were resuspended in Buffer H (25 mM HEPES pH 8.0, 2 mM MgCl2, 0.1 mM EDTA, 0.5 mM EGTA, 0.1% NP-40, 15% glycerol, 150 mM KCl) containing phosphatase inhibitors (0.1 mM Na-orthovanadate, 0.2 μM microcystin, 2 mM β-glycerophosphate, 1 mM Na pyrophosphate, 5 mM NaF) and protease inhibitors (20μg/mL leupeptin, 20μg/mL pepstatin A, 20μg/mL chymostatin, 200 μM phenylmethylsulfonyl fluoride). Lysates were ultracentrifuged at 98,500g for 90 min at 4°C. Protein G Dynabeads (Invitrogen) were conjugated with an α-FLAG antibody (Sigma) and immunoprecipitation of Dsn1-6His-3Flag was performed at 4°C for 3 hours. Beads were washed once with lysis buffer containing 2mM dithiothreitol (DTT) and protease inhibitors, 3 times with lysis buffer with protease inhibitors, 1 time in lysis buffer without inhibitors, and kinetochores particles were eluted by gentle agitation of beads in elution buffer (Buffer H + 0.5mg/mL 3FLAG Peptide (Sigma)) for 30 min at room temperature.

#### Immunoblot and silver stain analysis

Cell lysates were prepared as described above (kinetochore purification assay) or by bead-beat pulverization (Biospec Products) with glass beads in sodium dodecyl sulfate (SDS) buffer. Standard procedures for immunoblot and SDS-polyacrylamide gel electrophoresis (SDS-PAGE) were followed (Biggins et al., 1999). SDS-PAGE gels were transferred to 0.2μm nitrocellulose membrane (Bio-Rad) using either the semi-dry (Bio-Rad) or wet method (Hoefer). The anti-Mps1 antibodies were generated in rabbits against a recombinant Mps1 protein fragment (residues 440-764) of the protein by Genscript. The company provided affinity purified antibodies that we validated by purifying kinetochores from yeast strains with Mps1 or Mps1-13Myc and confirming that the antibody recognized a protein of the correct molecular weight that migrated more slowly with the 13Myc epitope tags. We subsequently used the antibody at a dilution of 1:10,000. The phospho-specific T74 Ndc80 antibody was generated by Pacific Immunology against the phospho-peptide spanning resides 69-80 of Ndc80 (KRTRS-pT-VAGGTN-Cys) and subsequently affinity purified. Immunoblotting with this antibody was performed at a dilution of 1:1,000 with 1μg/mL of non-phosphorylated competitor peptide (KRTRS-T-VAGGTN-Cys). The Ask1 antibody was affinity purified with recombinant GST-Ask1 from serum generated in rabbits against recombinant Dam1 complex and was used at a dilution of 1:5,000 (Gutierrez et al., 2020). The following commercial antibodies were used for immunoblotting: α-PGK1 (Invitrogen) 1:10,000, α-FLAG (M2; Sigma) 1:3,000, α-Myc (7D10; Cell Signaling) 1:1000, and α-HA (12CA5; Roche) 1:10,000 dilution. The α-Ndc80 antibody (OD4), used at 1:10,000, was a kind gift from Arshad Desai. Secondary antibodies used were donkey anti-rabbit antibody conjugated to horseradish peroxidase (HRP) (GE Biosciences) at 1:10,000 or a sheep anti-mouse antibody conjugated to HRP (Ge Biosciences) at 1:10,000. Antibodies were detected using SuperSignal West Dura Chemiluminescent Substrate (Thermo Scientific). Immunoblots were imaged with a ChemiDock MP system (Bio-Rad) or film. For silver stain analysis, protein samples were separated using pre-cast 4%-12% Bis-Tris Gels (Thermo-Fisher) and stained using the Silver Quest Staining Kit (Invitrogen).

#### Kinase assays

For radioactive kinase assays containing kinetochore particles (Figure 2A), kinetochores bound to beads were washed into kinase buffer (50 mM Tris-HCl pH 7.5, 75 mM NaCl, 5% glycerol, 10 mM MgCl_2,_ 1 mM DTT, 10 μM ATP) containing 66 nM γ-^32^P-ATP with either 5 µM reversine (Sigma R3904) or vehicle (DMSO) and incubated at 30 °C for 30 min. Kinetochores were eluted by boiling in sample buffer containing SDS and analyzed via silver stain and Phosphorimager. In addition to the previously detected phosphorylation on Spc105, Ndc80, and Dsn1 (London et al., 2012), we also detected phosphorylation on an additional unidentified protein (marked by an asterisk in Figure 2A), possibly due to altered SDS-PAGE conditions that allowed better resolution of kinetochore proteins. For kinase assays containing purified Ndc80 complex (Figure 5A), an anti-HA immunoprecipitation was performed and kinase assays were completed identically. Quantification of kinase assays was performed in Fiji (ImageJ) across three independent biological replicates. Intensity values from the Phosphorimager were normalized to Dsn1-His-Flag levels (Figure 2A) or Ndc80-3HA levels (Figure 5A) and the mean of the three replicates was used to calculate the change in phosphorylation between conditions.

#### Phosphatase assay

Native kinetochore particles were purified (see kinetochore purification assay) and kept bound to Dynabeads (Invitrogen). Beads were washed once with phosphatase buffer (Buffer H, 1mM MnCl), and then resuspended in phosphatase buffer with 200U lambda protein phosphatase (New England Biolabs) at 30 °C for 20 min. Control samples contained phosphatase inhibitors. Kinetochores were eluted by boiling beads in sample buffer containing SDS and phosphorylation was analyzed via immunoblotting.

### Biophysics

#### Laser trap instrument

The laser trap has been described previously (Franck et al., 2010). Position sensor response was mapped using the piezo stage to raster-scan a stuck bead through the beam, and trap stiffness was calibrated along the two principle axes using the drag force, equipartition, and power spectrum methods. Force feedback was implemented with custom LabView software. During force measurements, bead-trap separation was sampled at 40 kHz while stage position was updated at 50 Hz to maintain the desired tension (force-clamp assay) or ramp-rate (force-ramp assay). Bead and stage position data were decimated to 200 Hz before storing to disk.

#### Bead functionalization and slide preparation for laser trap experiments

Native kinetochore particles were linked to beads as previously described (Franck et al., 2010). First, streptavidin-coated polystyrene beads (0.56 μm in diameter, Spherotech Inc., Libertyville IL) were functionalized with biotinylated anti-5His antibodies (Qiagen, Valencia CA) and stored with continuous rotation at 4°C in BRB80 (80 mM PIPES, 1 mM MgCl2, and 1 mM EGTA, pH 6.9) supplemented with 8 mg·mL^-1^ BSA for up to 3 months. Immediately prior to each experiment, beads were decorated with kinetochore particles by incubating 6 pM anti-5His beads for 60 min at 4°C with purified kinetochore material, corresponding to Dsn1-His-Flag concentrations ranging between from 2 to 4 nM. The resulting fraction of active beads capable of binding microtubules remained below 50%, thus ensuring single particle conditions (Akiyoshi et al., 2010; Sarangapani et al., 2013; Sarangapani et al., 2014).

Flow chambers (∼10 µL volume) were made using glass slides, double-stick tape, and KOH-cleaned coverslips, and then functionalized in the following manner. First, 15 to 20 µL of 10 mg·mL^−1^ biotinylated BSA (Vector Laboratories, Burlingame CA) was introduced and allowed to bind to the glass surface for 15 min at room temperature. The chamber was then washed with 100 µL of BRB80. Next, 25 µL of 1 mg·mL^−1^ avidin DN (Vector Laboratories, Burlingame CA) was introduced, incubated for 3 min, and washed out with 100 µL of BRB80. GMPCPP-stabilized biotinylated microtubule seeds were introduced in BRB80, and allowed to bind to the functionalized glass surface for 3 min. The chamber was then washed with 100 µL of growth buffer (BRB80 containing 1 mM GTP and 1 mg·mL^−1^ κ-casein). Finally, kinetochore particle-decorated beads were introduced at eight to ten-fold dilution from the incubation mix (see above) in a solution of growth buffer containing 1.5 mg·mL^−1^ purified bovine brain tubulin and an oxygen scavenging system (1 mM DTT, 500 µg·mL^−1^ glucose oxidase, 60 µg·mL^−1^ catalase, and 25 mM glucose).

For the ATP (or mock) exposure experiments (Figures 1, 2, 3 and 5; Supplemental Figures S1 and S2), 6 mM ATP or 6 mM ADP (conjugated with sodium salt; Sigma-Aldrich, St. Louis MO) were included along with the contents described above. In the case of phosphatase add-back experiments (Figure 1), 1200 units of lambda phosphatase were added to each slide. For the reversine add-back experiments (Figure 2), 5 µM reversine suspended in DMSO was included along with the contents described above. An equivalent volume of DMSO was used in the corresponding non-reversine control experiments.

The edges of the flow chamber were sealed to prevent evaporation, and the time was set to 0 min. All laser trap experiments were performed in temperature-controlled rooms, maintained at 23 °C, for up to 90 min after the chamber was sealed.

#### Rupture force measurements

For efficiency of data collection, beads that were already bound to microtubules (on the lattice, away from the dynamic tip) were usually chosen for measurements of rupture strength. Initially, the attachments were preloaded with a constant tensile force of 1 to 3 pN, which caused the lattice-bound beads to slide until reaching the microtubule plus end. Once at the end, we verified that the beads moved under the preload force at a rate consistent with that of microtubule growth or shortening. The laser trap was subsequently programmed to ramp the force at a constant rate (0.25 pN·s^-1^) until the linkage ruptured, or until the load limit of the trap was reached (∼23 pN under the conditions used here). Fewer than 5% of all trials ended in detachment during the preload period before force ramping began, while 0 to 15% reached the load limit (depending on the conditions tested). These out-of-range events were included in the median force calculations and the survival probability distributions. Under selected conditions, we also tested beads that were floating freely in solution in order to estimate the fraction of active beads, capable of binding microtubules. When beads decorated with wild type kinetochore particles (from SBY8253) were tested in the absence of adenosine, six beads out of thirty tested (20%) bound to microtubules. Similarly, when identically prepared beads were tested in the presence of ADP, five beads out of twenty two (23%) bound microtubules. However, when the beads were tested in the presence of ATP, only three out of twenty seven tested (11%) bound to microtubules, suggesting that the binding activity of the kinetochore particles (like their rupture strength) is reduced upon exposure to ATP. We found no statistically significant difference in the rupture strengths of pre-bound versus free beads. Most of the measurements (> 98%) were obtained using pre-bound bead-microtubule pairs.

Supplemental Table S4 summarizes the median rupture force values for each kinetochore type or condition. All the individual rupture force values, the Kaplan-Meier survival probability estimates (with 95% confidence intervals), the numbers of trials that reached the load limit of the trap without rupture, and other statistics (e.g., p-values from log-rank tests), are included in Supplemental Table S5.

---

**Supplemental Figure S1.**
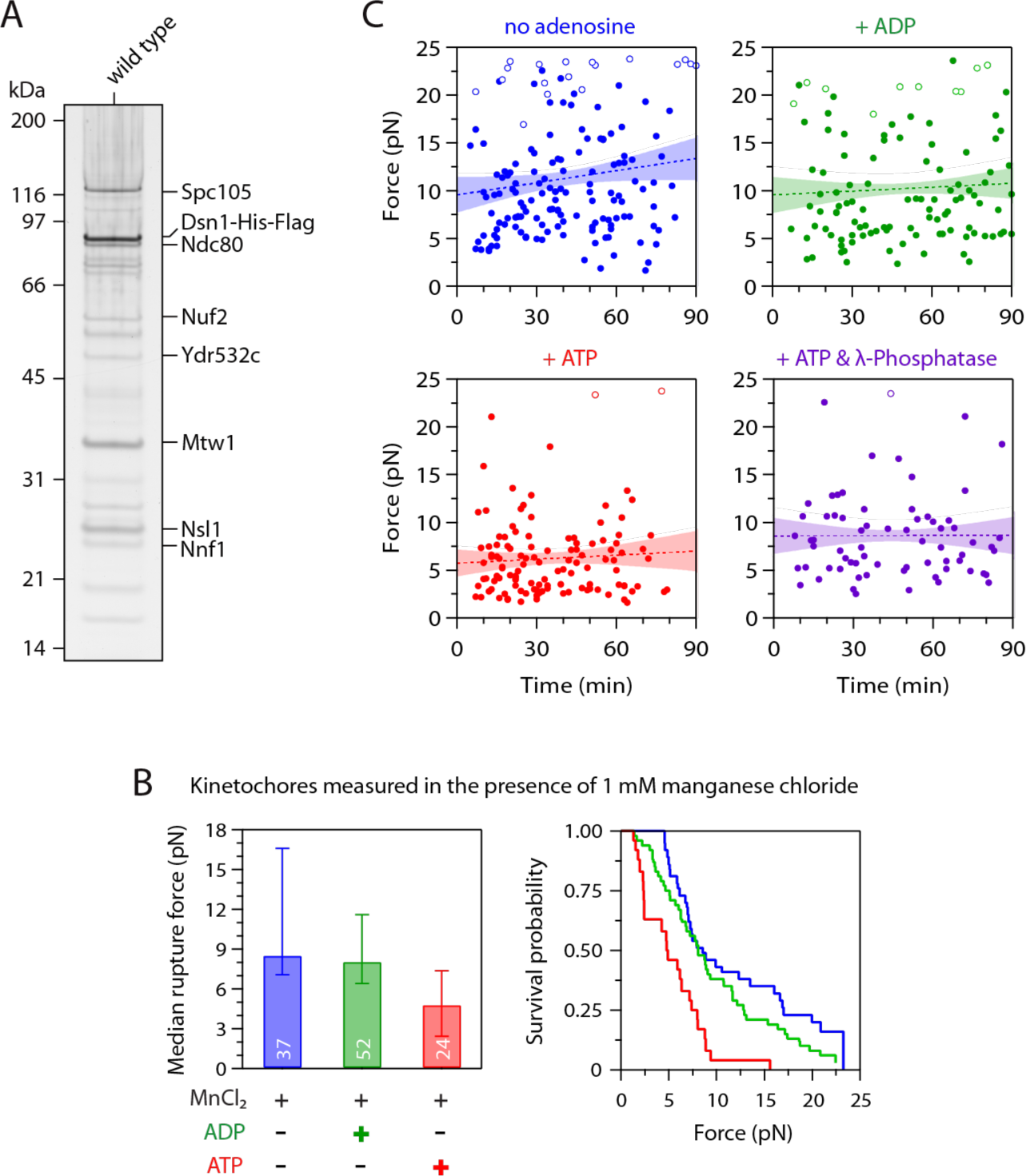
Control experiments related to Figure 1. **(A)** Kinetochore material purified by immunoprecipitation of Dsnl-His-Flag from wild type cells (SBY8253) visualized by silver stain after SDS-PAGE. **(B)** Control experiments related to Figure 1D. Median rupture strengths (left) and corresponding survival probability distributions (right) for wild type kinetochores (from SBY8253) in the presence of 1 mM manganese chloride (MnCI,). The kinetochores under these conditions were weakened upon exposure to ATP by an a mount similar to that in the absence of manganese chloride. Values inside bars indicate numbers of events for each condition. Error bars represent ± 95% confidence intervals calculated by bootstrapping (see Supplemental Material). (**C**) The ATP-dependent weakening reaction is completed within minutes. Individual rupture force values (solid circles) are plotted against the time elapsed since the kinetochore-decorated beads were prepared as indicated (without adenosine, or mixed with ATP, ADP, or ATP & α-phosphatase). Right-censored events that reached the load limit of the laser trap before rupture are also plotted (open circles). The slopes of lines fit to all the data (dotted lines, with 95% confidence intervals shown) were not significantly different from zero (0.42 ± 0.45 pN min^-1^ 0.014 ± 0.045 pN min^-1^ 0.014 ± 0.040 pN min^-1^ and 0.000 ± 0.049 pN min^-1^ respectively), indicating that ATP-dependent weakening occurred during the ~10 min slide preparation, with no significa nt weakening thereafter.

**Supplemental Figure S2.**
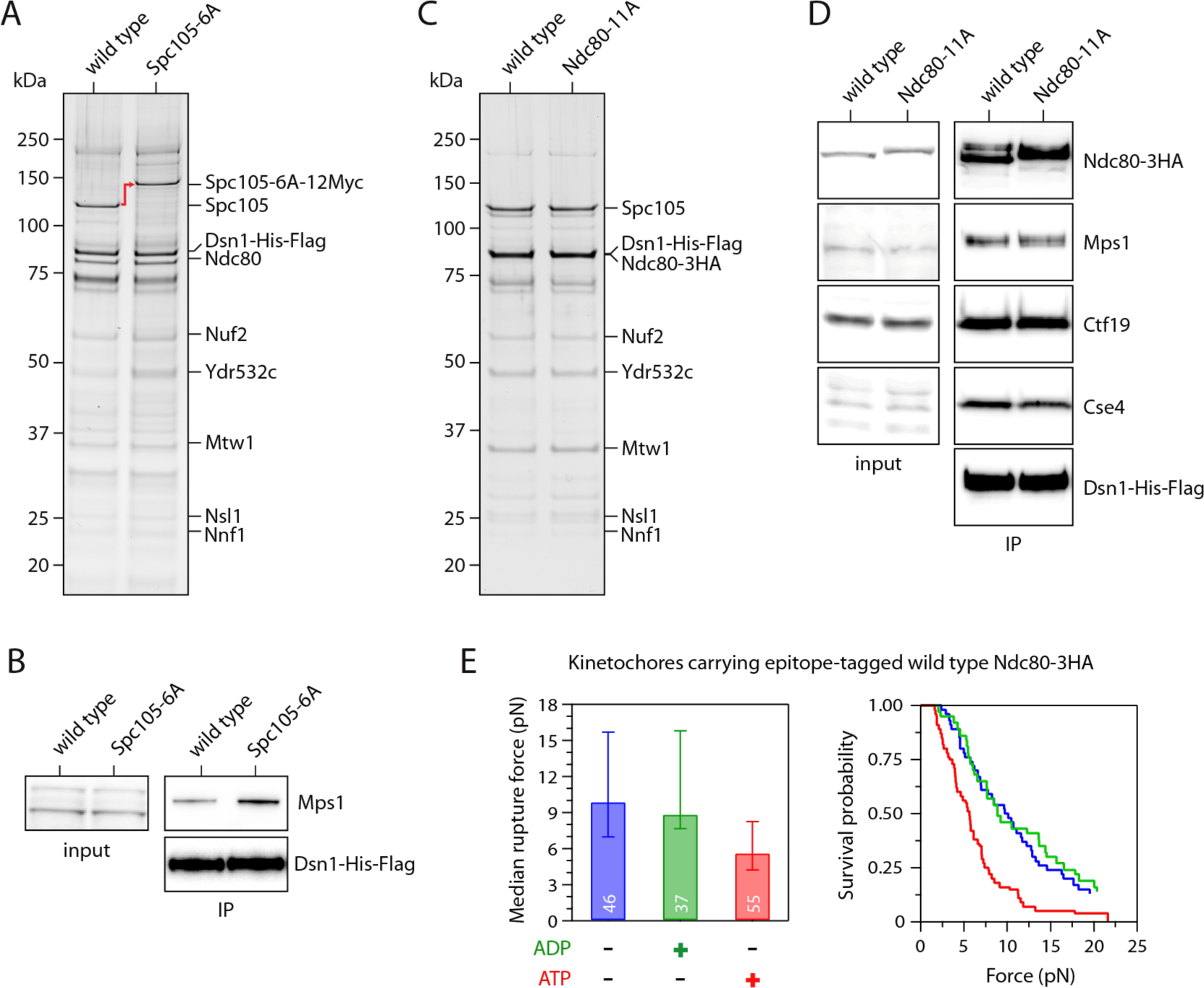
Control experiments related to Figure 3. **(A)** Kinetochore material purified by immunoprecipitation of Dsn 1-His-Flag from mutant cells carrying phospho-deficient Spcl 05-6A-12Myc (SBY10315) and from wild type cells (SBY8253) visualized by silver stain after SDS-PAGE. **(B)** Kinetochores purified by immunoprecipitation of Dsn 1-His-Flag from cells carrying epitope-tagged wild type Spcl 05-12Myc (SBY10336) and from mutant cells carrying phospho-deficient Spcl 05-6A-12Myc (SBY10315) were analyzed by immunoblotting for Mpsl levels. No Dsn 1-His-Flag blot is shown for the input material because Dsn 1 was undetectable in whole cell lysates, due to an overlapping cross-reactive band. (C) Kinetochore material purified by immunoprecipitation of Dsn 1-His-Flag from cells carrying epitope-tagged wild type Ndc80-3HA (SBYl 1808) and from mutant cells carrying phospho-deficient Ndc80-11A-3HA (SBY19380) visualized by silver stain after SDS-PAGE. **(D)** Similar levels of Mpsl copurify with wild type and Ndc80-11A kinetochores. Kinetochore material was purified by immunoprecipitation of Dsnl-Hls-Flag (from SBYl 1808 and SBY19380) and analyzed by immunoblotting. No Dsnl-His-Flag blot is shown for the input material because Dsn 1 was undetectable in whole cell lysates, due to an overlapping cross-reactive band. **(E)** Control experiments related to Figure 3D. Median rupture strengths (left) and corresponding survival probability distributions (right) for kinetochores carrying epitope-tagged wild type Ndc80-3HA (from SBYl 1808). These kinetochores were weakened upon exposure to ATP by an amount similar to kinetochores carrying untagged wild type Ndc80. Values inside bars indicate numbers of events for each condition. Error bars represent ± 95% confidence intervals calculated by bootstrapping (see Supplemental Material).

**Supplemental Figure S3.**
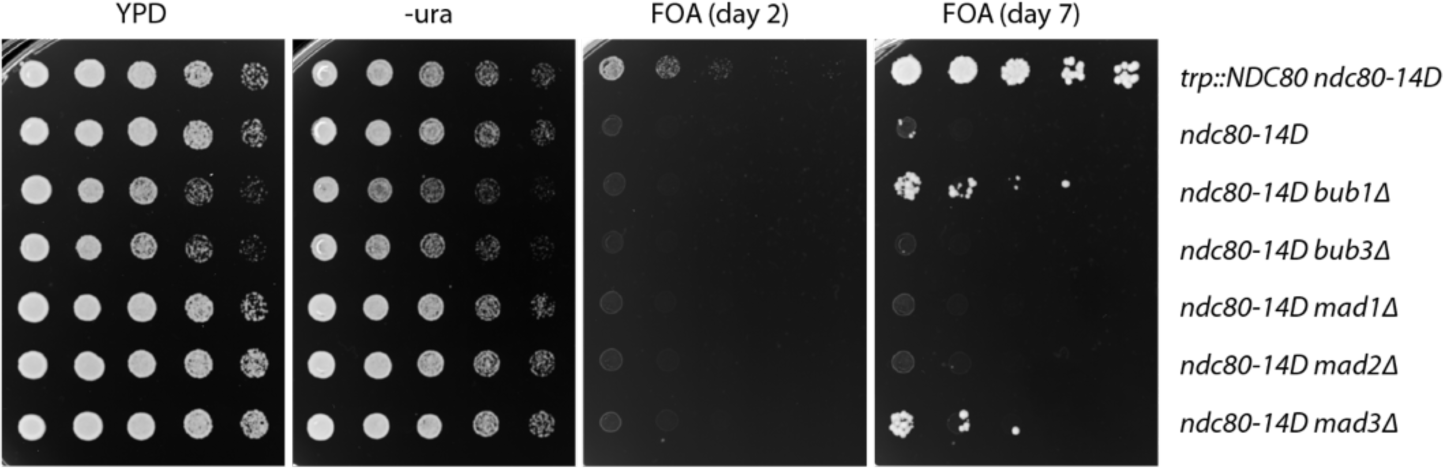
Control experiments related to Figure 4. The phospho-mimetic *ndc80-14D* allele is lethal and its lethality is not rescued by deletion of the checkpoint gene *MAD2*. However, suppressors do arise after many days of growth. The indicated strains (SBYl 5149, SBY9453, SBYl 5139, SBYl 7991, SBYl 7993, SBYl 5087, SBYl 1334) were tested using a plasmid shuffle assay,where growth on 5′fluorouracil (FOA) plates indicates viabil ity in the absence of a covering copy of wild type *NDC80*. Five-fold serial dilutions were plated onto YPD, -ura, or FOA plates and grown at 23•C for two days. After seven days of growth on FOA, a small number of colonies grow from the *ndc80-14D* (SBY9453), *ndc80-14D bub1*Δ (SBYl 5139), and *ndc80-14D mad3*Δ (SBYl1334) strains.

**Supplemental Figure S4.**
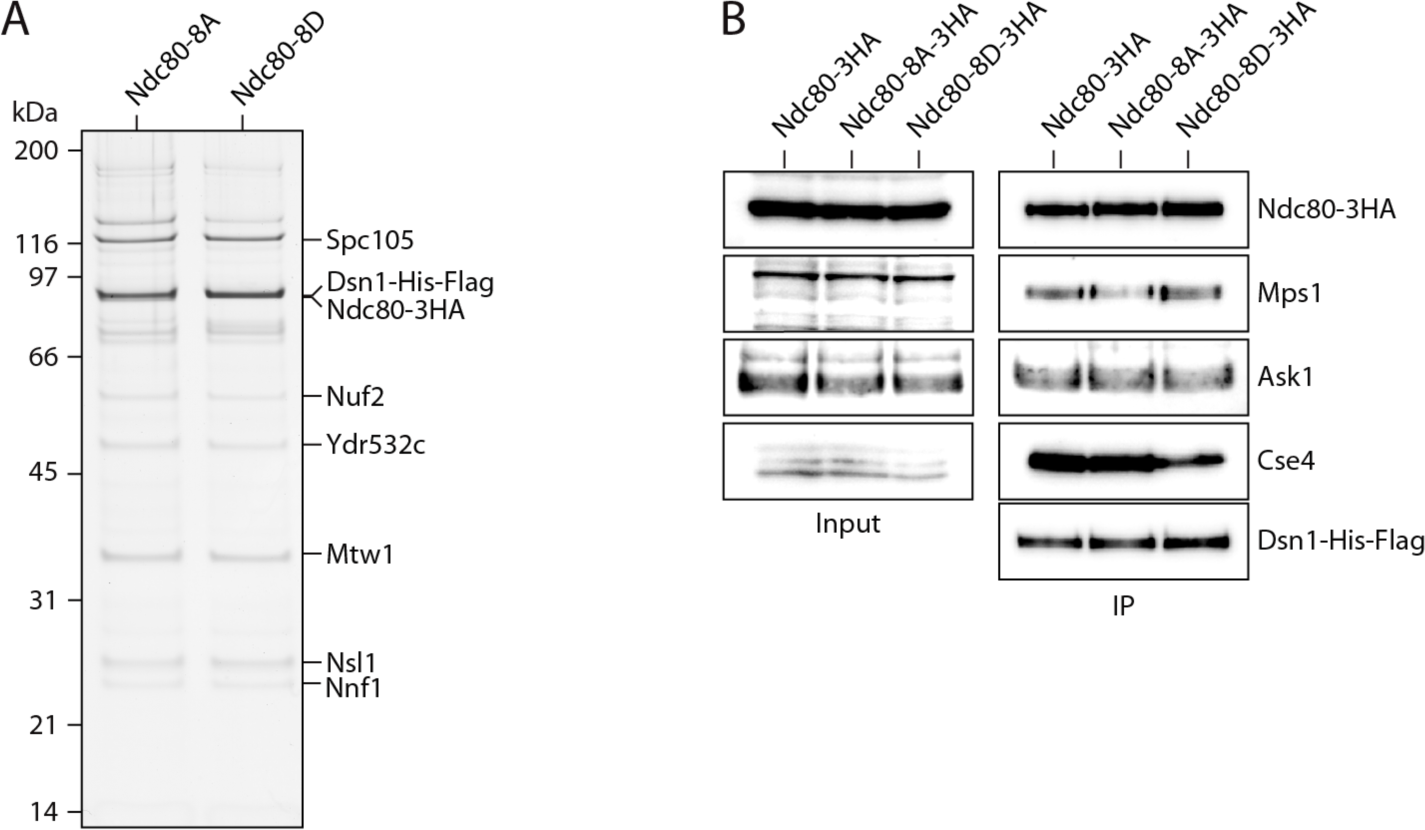
Control experiments related to Figure 5. **(A)** Kinetochore material purified by immunoprecipitation of Dsnl-His-Flag from cells carrying phospho-deficient Ndc80-8A (SBY19855) and cells carrying phospho-mimetic Ndc80-8D (SBY19877) visualized by silver stain after SDS-PAGE. **(B)** Kinetochore material purified by immunoprecipitation of Dsn1-His-Flag from Ndc80-3HA (SBY19186), Ndc80-8A-3HA (SBY19855), and Ndc80-8D-3HA (SBY19877) was analyzed for kinetochore protein composition via immunoblotting. No Dsnl-His-Flag blot is shown for the input material because Dsnl was undetectable in whole cell lysates, due to an overlapping cross-reactive band.

**Supplementary Table 1.**
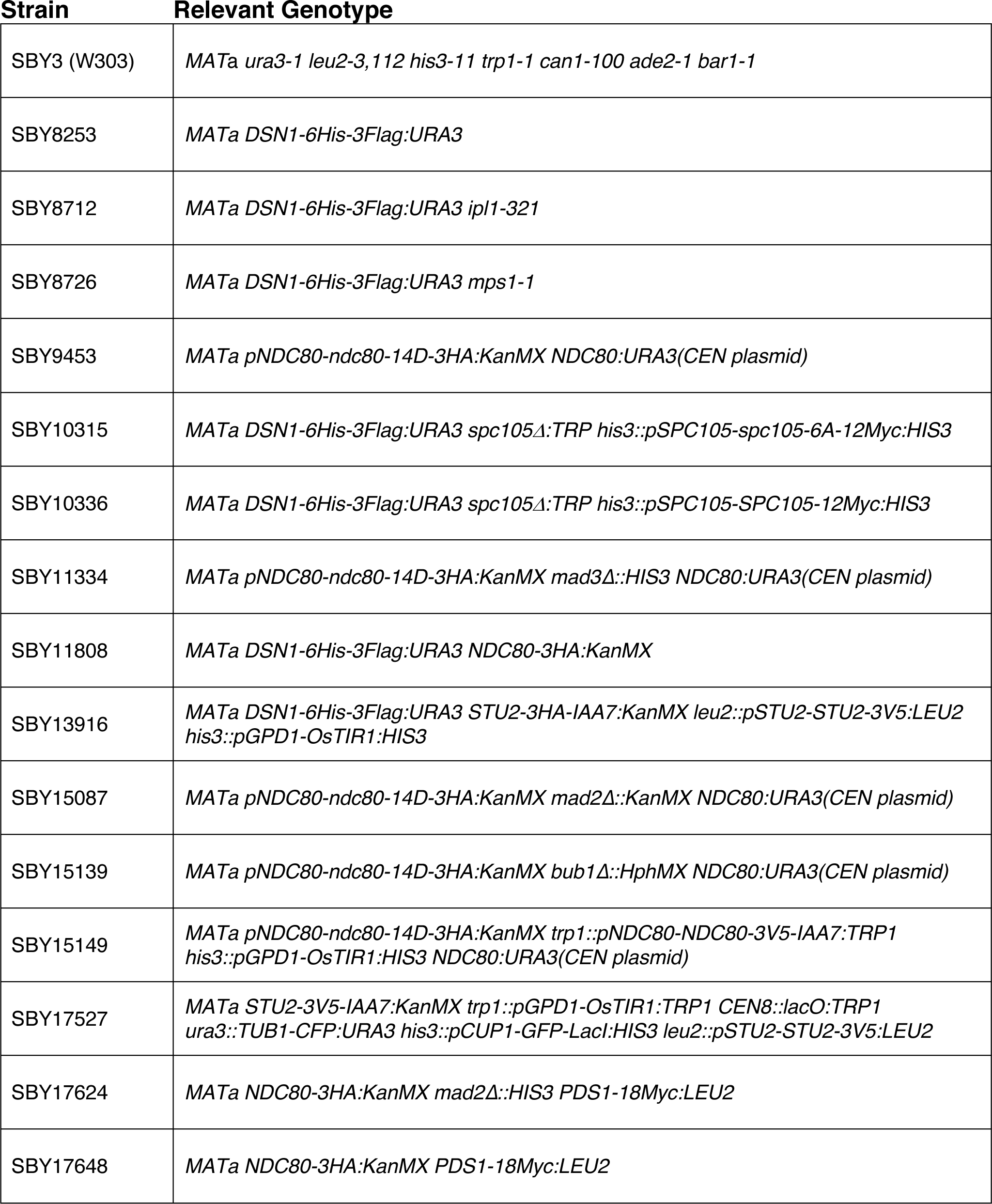
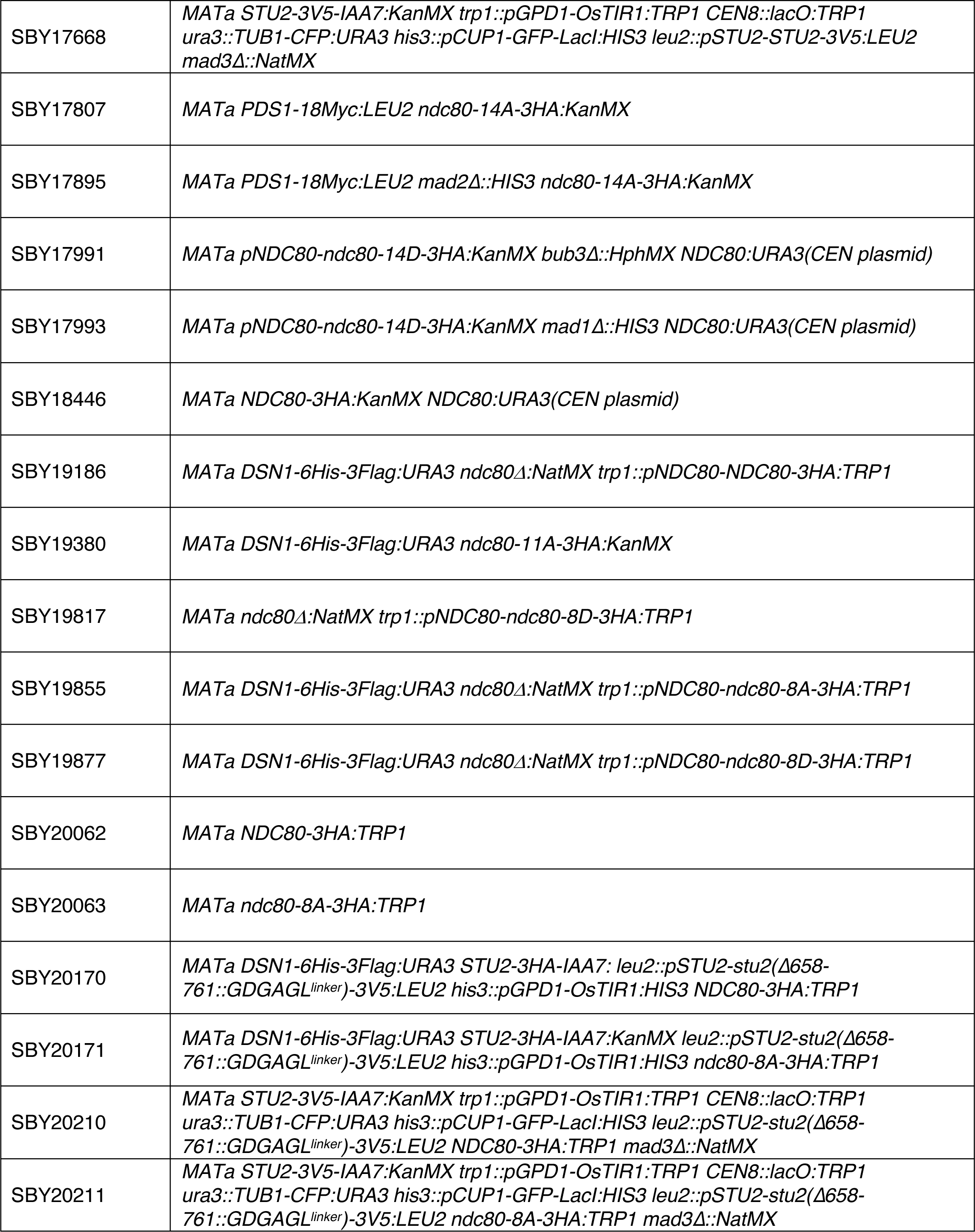
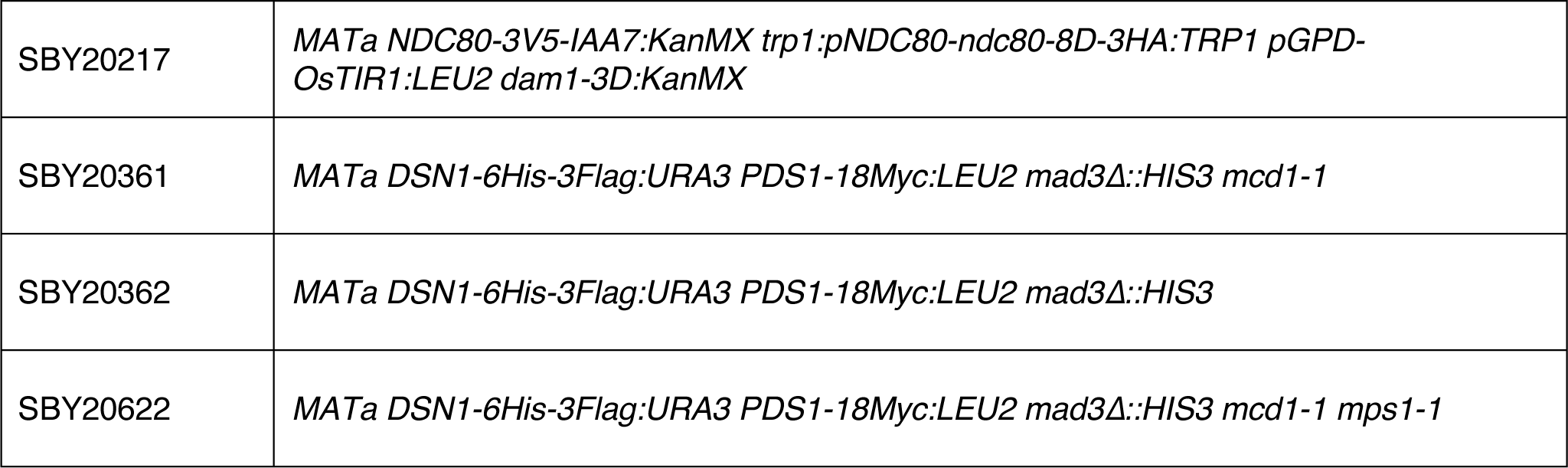
Strains used in this study. All strains are derivatives of SBY3 (W303)

**Supplementary Table 2.** Plasmids used in this study

| Plasmid | Description | Mutations |
| --- | --- | --- |
| pSB1302 | *pNDC80-NDC80:URA3* (CEN plasmid) | none |
| pSB1577 | *pNDC80-ndc80-14D:KanMX6* (pSK981, gift from Lechner lab) | S4D, T5D, S6D, T21D, S22D, S37D, T38D, T43D, T74D, T79D, T82D, S205D, T248D, T252D |
| pSB1848 | *pNDC80-ndc80-14A:KanMX6* (pSK1039, gift from Lechner lab) | S4A, T5A, S6A, T21A, S22A, S37A, T38A, T43A, T74A, T79A, T82A, S205A, T248A, T252A |
| pSB2412 | *pNDC80-NDC80-3HA:TRP1* | none |
| pSB3131 | *pNDC80-ndc80-11A-3HA:KanMX6* (derivative of pSB1848) | S4A, T5A, S6A, S22A, T38A, T43A, T79A, T82A, S205A, T248A, T252A |
| pSB3207 | *pNDC80-ndc80-8A-3HA:TRP1* (derivative of pSB2412) | S4A, T5A, S6A, S22A, T38A, T43A, T79A, T82A |
| pSB3208 | *pNDC80-ndc80-8D-3HA:TRP1* (derivative of pSB2412) | S4D, T5D, S6D, S22D, T38D, T43D, T79D, T82D |

**Supplementary Table 3.**
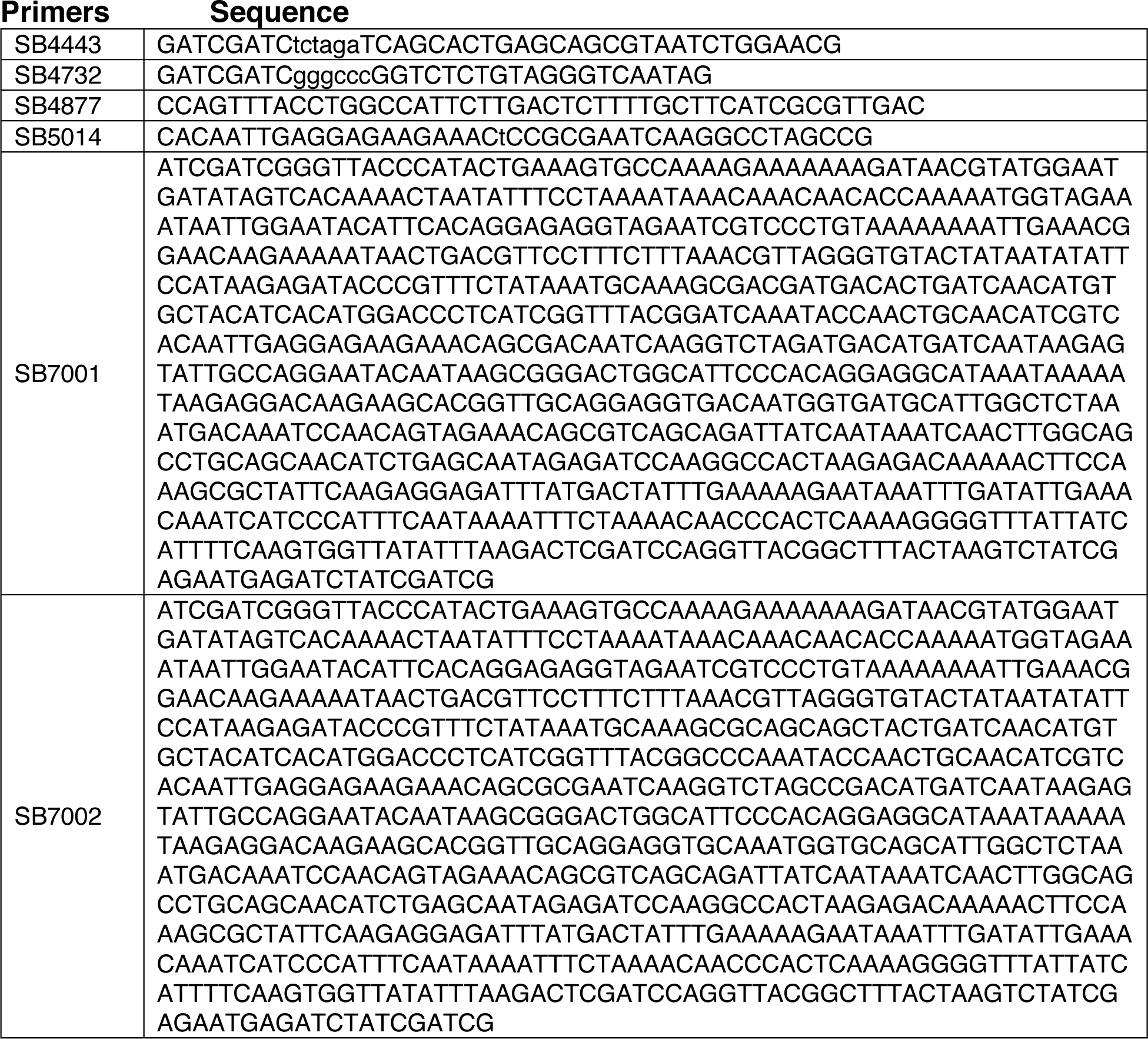
Primers used in this study (all listed 5′ to 3′)

**Supplemental Table S4.** Summary of laser trap results. Rupture forces indicate median ± σ obtained from bootstrapping of N individual rupture events, with replacement. All the individual rupture force values are provided in Supplemental Table S5.

| Figure 1 & S1 | Strain Number | Kinetochores Type, Treatment | [Dsn1] (nM) | Median Rupture Force (pN) (N) |
| --- | --- | --- | --- | --- |
| | SBY8253 | wild type, no treatment | 2 | 9.83 ± 0.50 (156) |
| | | wild type, ADP-treated | 2 | 8.66 ± 0.56 (110) |
| | | wild type, ATP-treated | 2 | 5.31 ± 0.56 (118) |
| | | wild type, ATP- & Phosphatase-treated | 2 | 7.53 ± 0.62 (69) |
| | | wild type, MnCl<sub>2</sub>-treated | 2 | 8.57 ± 1.45 (37) |
| | | wild type, MnCl<sub>2</sub>- & ADP-treated | 2 | 8.08 ± 0.84 (52) |
| | | wild type, MnCl<sub>2</sub>- & ATP-treated | 2 | 4.84 ± 1.12 (24) |

| Figure 2 | Strain Number | Kinetochores Type, Treatment | [Dsn1] (nM) | Median Rupture Force (pN) (N) |
| --- | --- | --- | --- | --- |
| | SBY8253 | wild type, DMSO-treated | 2 | 8.15 ± 0.64 (85) |
| | | wild type, DMSO- & ADP-treated | 2 | 8.20 ± 0.44 (80) |
| | | wild type, DMSO- & ATP-treated | 2 | 4.97 ± 0.39 (100) |
| | | wild type, DMSO- & Reversine-treated | 2 | 8.11 ± 0.66 (74) |

| Figure 3 & S2 | Strain Number | Kinetochores Type, Treatment | [Dsn1] (nM) | Median Rupture Force (pN) (N) |
| --- | --- | --- | --- | --- |
| | SBY10315 | Spc105-6A (Myc), no treatment | 4 | 8.34 ± 0.38 (84) |
| | | Spc106-6A (Myc), ADP-treated | 4 | 8.66 ± 0.60 (57) |
| | | Spc105-6A (Myc), ATP-treated | 4 | 3.17 ± 0.25 (59) |
| | SBY19380 | Ndc80-11A (3HA), no treatment | 3 | 8.66 ± 0.64 (96) |
| | | Ndc80-11A (3HA), ADP-treated | 3 | 10.04 ± 0.98 (74) |
| | | Ndc80-11A (3HA), ATP-treated | 3 | 8.53 ± 0.62 (71) |
| | SBY11808 | Ndc80 (3HA), no treatment | 3 | 9.92 ± 1.63 (46) |
| | | Ndc80 (3HA), ADP-treated | 3 | 8.87 ± 2.46 (37) |
| | | Ndc80 (3HA), ATP-treated | 3 | 5.66 ± 0.46 (55) |

| Figure 5 | Strain Number | Kinetochores Type, Treatment | [Dsn1] (nM) | Median Rupture Force (pN) (N) |
| --- | --- | --- | --- | --- |
| | SBY19855 | Ndc80-8A (3HA), no treatment | 4 | 9.36 ± 0.71 (111) |
| | | Ndc80-8A (3HA), ADP-treated | 4 | 8.64 ± 0.54 (79) |
| | | Ndc80-8A (3HA), ATP-treated | 4 | 9.41 ± 0.48 (132) |
| | SBY19877 | Ndc80-8D (3HA), no treatment | 4 | 5.60 ± 0.37 (135) |

